# A CUG-initiated CATSPERθ functions in the CatSper channel assembly and serves as a checkpoint for flagellar trafficking

**DOI:** 10.1101/2023.03.17.532952

**Authors:** Xiaofang Huang, Haruhiko Miyata, Huafeng Wang, Giulia Mori, Rie Iida-Norita, Masahito Ikawa, Riccardo Percudani, Jean-Ju Chung

## Abstract

Calcium signaling is critical for successful fertilization. In spermatozoa, calcium influx into the sperm flagella mediated by the sperm specific CatSper calcium channel is necessary for hyperactivated motility and male fertility. CatSper is a macromolecular complex and is repeatedly arranged in zigzag rows within four linear nanodomains along the sperm flagella. Here, we report that the *Tmem249*-encoded transmembrane domain containing protein, CATSPERθ, is essential for the CatSper channel assembly during sperm tail formation. CATSPERθ facilitates the channel assembly by serving as a scaffold for a pore forming subunit CATSPER4. CATSPERθ is specifically localized at the interface of a CatSper dimer and can self-interact, suggesting its potential role in CatSper dimer formation. Male mice lacking CATSPERθ are infertile because the sperm lack the entire CatSper channel from sperm flagella, rendering sperm unable to hyperactivate, regardless of their normal expression in the testis. In contrast, genetic abrogation of any of the other CatSper transmembrane subunits results in loss of CATSPERθ protein in the spermatid cells during spermatogenesis. CATSPERθ might acts as a checkpoint for the properly assembled CatSper channel complex to traffic to sperm flagella. This study provides insights into the CatSper channel assembly and elucidates the physiological role of CATSPERθ in sperm motility and male fertility.

## Introduction

Ion channels are transmembrane (TM) proteins that form an aqueous pore by which certain ions can pass across the plasma membrane. Sensing chemical and electrical stimuli, ion channels open and redistribute ions, which can affect various cellular processes from electrical excitation of neuron, muscle, and heart to locomotion (1-4). To fine-tune these critical functions, many ion channels form macromolecular complexes that contain not only pore-forming α-subunits but also other ancillary subunits (1). For example, the α-subunits of the voltage-gated L-type calcium channel (Ca_v_ 1.1-1.4) co-assemble with α2δ, β and/or γ subunits (5, 6). The homotetrameric α-subunits of the voltage-gated potassium channel K_v_4.2 can further complex with KChIP1 and DPP6S as dodecamer (7, 8). Proper assembly of such ion channel complexes is crucial for cellular homeostasis and strictly regulated to achieve appropriate channel density and function in the plasma membrane (1). For example, the chaperone proteins DNAJB12 and DNAJB14 regulate the tetrameric assembly of ERG (ether-a-go-go-related) type K^+^ channel (9). The intracellular N-terminal T1 domain regulates the tetramerization of the K_v_ channel Shaker (10, 11). Failure of such assembly often leads to inappropriate targeting and/or aberrant function of the channel complex, resulting in cellular pathophysiology (12-14).

In mammalian spermatozoa, the CatSper calcium channel, which mediates calcium influx to develop the hyperactivated motility required for successful fertilization (15), is the best example of macromolecular ion channel complex. Previous biochemical and structural studies have revealed that the CatSper channel is composed of the pore-forming subunits CATSPER1, 2, 3 and 4 (16-18), TM auxiliary subunits including CATSPERβ, γ, δ, ε, η, TMEM249 and an unannotated single TM protein, cytosolic subunits containing the subcomplex EFCAB9-CATSPERζ and CATSPERτ (19-25), and an uncharacterized transporter, SLCO6C1 (19, 24, 26). Unlike other ion channels, the CatSper complexes are uniquely arranged along the sperm flagellum, forming four linear Ca^2+^ signaling nanodomains (27). Our recent 3D reconstructed cryo-tomograms of mouse sperm flagella revealed that the CatSper complexes are arranged in zigzag rows in each quadrant from extracellular views (26). Intracellular views further visualized diagonal arrays below two staggered adjacent channel complexes, suggesting that CatSper forms dimers that are building blocks for the zigzag rows. Male mice lacking TM subunits, either any of the pore-forming CATSPER1-4 subunits or the auxiliary TM subunit CATSPERδ, are infertile due to the absence of the entire complex in the sperm flagella (16, 17, 23, 28). These results suggest that only correctly assembled, complete CatSper channel complex can be trafficked to the flagellum. Moreover, the integrity of the CatSper nanodomains and the continuity of the zigzag rows are associated with the ability to properly develop hyperactivated motility and the chirality of the sperm swimming path (22, 24, 26, 29). The sperm that successfully arrived at the fertilization site are recognized with the intact CatSper nanodomains as a molecular signature (30). Despite the physiological significance of the CatSper nanodomains and their underlying higher-order arrangement, it remains unknown what controls the formation of monomers and dimers of the CatSper channel complex and the molecular mechanisms that link the complex assembly to complex trafficking.

Here, we report that *Tmem249* has coevolved with other known CatSper genes and uniquely uses a CTG start codon to encode CATSPERθ. Genetic ablation of *Catsperq* in mice results in the incompletely assembled CatSper complexes that are unable to target to the sperm flagella, rendering male mice infertile due to defective sperm hyperactivation. We found that CATSPERθ not only stabilizes the CATSPER4 subunit to form the monomeric CatSper complex, but also self-interacts to contribute to the dimer formation required for the higher order zig-zag arrangement of the channels. Our studies provide molecular insights into the assembly mechanisms of the CatSper channel complex and its crosstalk in complex trafficking during sperm tail formation.

## Results

### A comprehensive genomic screening identifies *Tmem249* in the CatSper coevolutionary gene network

Most of the genes encoding the known CatSper subunits share three common features: they are specifically expressed in the testis, they show similar patterns of lineage-specific gain and loss during evolution, and the encoded proteins display interdependency of protein levels in the spermatozoa (24). Taking advantage of this gene coevolution, we performed a comprehensive genomic screen to search for genes that can be clustered together with the CatSper genes. We obtained the CatSper coevolutionary network by automatic clustering of significant pairwise associations among 60,675 orthologous gene groups from 1,256 eukaryotic genomes. A previously uncharacterized *Tmem249* gene clustered together with the coevolutionary network of known CatSper genes (*Catsper1-4, Catsperb-d*, and *Efcab9*) (**Fig. 1** and **Fig. S1*A***). Reciprocally, CatSper genes constitute eight of the ten most significantly coevolving genes of *Tmem249* (**Fig. S1*B***). Consistent with the mRNA expression patterns of other CatSper genes, *Tmem249* is also specifically expressed in the testis (**Fig. S1*C***), meeting two of the criteria for CatSper genes. During the preparation of this manuscript, an independent study also reported that TMEM249 is likely a component in the purified mouse CatSper complex and modeled it in the single particle cryo-EM structure of the CatSper channel complex (19), further supporting TMEM249 as a strong candidate for a new CatSper component.

**Fig. 1.**
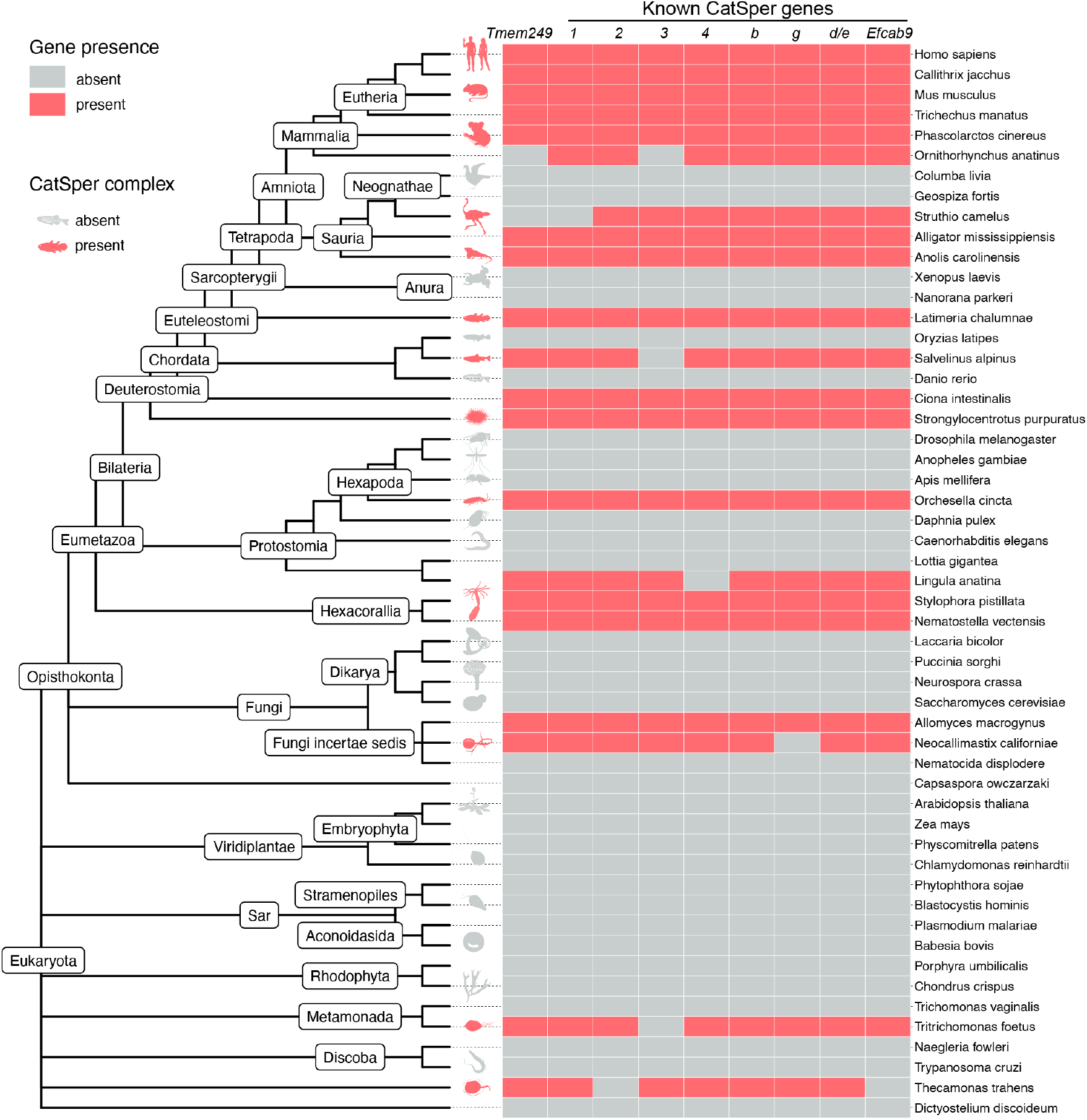
*Tmem249* is co-evolving with other CatSper genes. Distribution map of *Tmem249* and other CatSper genes across eukaryotes. The phylogenetic tree and gene classification is according to NCBI taxonomy and OrthoDB, respectively. Shown are 53 representative species of the full eukaryotic dataset (1,256 species) analyzed in this study.

### TMEM249 protein levels and localization are dependent on the CatSper channel in the sperm flagella

We first investigated whether the absence of the CatSper channel affects TMEM249 protein level and localization as shown in other known CatSper subunits. Confocal imaging of immunostained TMEM249 in *wt* and *Catsper1*^*-/-*^ mouse sperm from the cauda epididymis shows that it localizes specifically and CatSper-dependently to the principal piece like CATSPER1 (**Fig. 2*A***). Super resolution imaging by 3D Structured Illumination Microscopy imaging (SIM) further reveals that TMEM249 displays 4-linear arrangement along the sperm tail (**Fig. 2*B***), a distinct pattern previously observed for CatSper nanodomains (22, 24, 25, 27). Immunoblot analysis further confirms that TMEM249 is an integral part of the CatSper complex unit; it is not detected in *Catsper1*^*-/-*^ or *Catsperd*^*-/-*^ sperm in which the entire channel is absent but is detected to a much lesser extent in *Efcab9*^*-/-*^ and *Catspert*^*-/-*^ sperm (**Fig. 2*C-D***). All these results suggest that TMEM249 is associated with the CatSper complex in cauda sperm. We herein name it CATSPERθ.

**Fig. 2.**
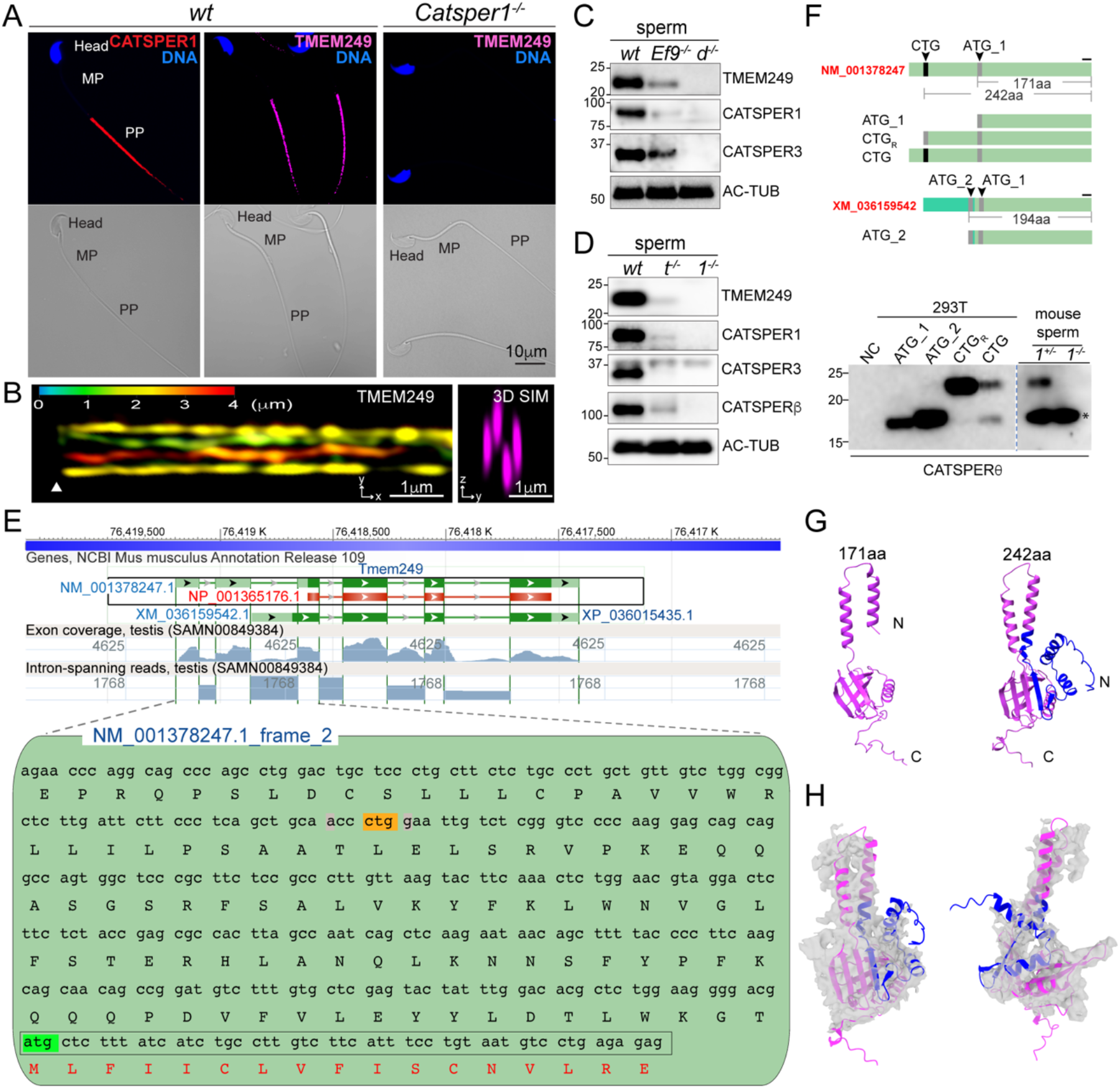
CTG initiated TMEM249 protein level in sperm are CatSper dependent. (*A*) Immunofluorescence confocal images (*upper*) of TMEM249 and CATSPER1 in mouse epididymal sperm. TMEM249 is detected in the principal piece (PP) of *wt* mouse sperm (*middle*, magenta), consistent with that of CATSPER1 (*left*, red), but not in *Catsper1*^*-/-*^ sperm (right). Sperm head is shown by counterstaining DNA with Hoechst (blue). The corresponding differential interference contrast (DIC) to the fluorescent image are shown (*bottom*). MP, midpiece. (*B*) 3D structural illumination microscope (SIM) imaging of TMEM249 in *wt* mouse sperm. *x-y* projection (*left*) and *y-z* projection (*right*) are shown; *z* axis information is color-coded in *x-y* projection. Arrowheads indicate the annulus. (*C*-*D*) Immunoblot analysis of TMEM249 and other CatSper subunits in *wt, Efcab9*^*-/-*^ (*Ef9*^*-/-*^), *Catsperd*^*-/-*^ (*d*^*-/-*^) (*C*), *Catspert*^*-/-*^ (*t*^*-/-*^) and *Catsper1*^*-/-*^ (*1*^*-/-*^) sperm (*D*). Acetylated tubulin (AC-TUB) is probed as loading control. (*E*) NCBI sequence-viewer representation of the *Mus musculus Tmem249* (*Catsperq*) locus (*upper*) and hypothetical translation of its 5’-UTR of (*lower*). Exons are represented by green boxes in which the coding sequence are in dark green. The reference mRNA (NM_001378247.1) and protein (NP_001365176.1) principal variants are framed in black box. RNA-seq exon coverage and intron spanning reads for the testis dataset (SAMN00849384) are shaded in blue (linear scale). The ATG-initiated ORF (boxed in gray, ATG highlighted in green) is in frame with candidate initiation codon CTG (in yellow) in the 5’-UTR. Nucleotides at positions -3 and +4 (highlighted in gray) from the CTG match to preferred Kozak consensus. (*F*) Schematic illustration of the nucleotide sequences used to express *Mus musculus* CATSPERθ proteins in 293T cells (*upper*). ATG is in grey, CTG is in black. The N-terminal sequence of XM_036159542 (boxed in fluorescent green) is different from that of NM_001378247. ATG_1 (NP_001365176.1, 171 aa), ATG_2 (XP_036015435, 194 aa), CTG_R_ (ORF start from CTG but with CTG replaced with ATG, 242 aa) and CTG (ORF start from CTG in the native 5’-UTR). The black line above the C-terminal sequence common to all variants indicates the location of the corresponding antigen of CATSPERθ antibody used in this study. The yield protein product is compared with that in native *Catsper1*^*+/-*^ (*1*^*+/-*^) and *Catsper1*^*-/-*^ (*1*^*-/-*^) sperm by western blot analysis (*lower*). The dotted line indicates different exposure time of the same membrane. Asterisk in the mouse sperm blot indicates the non-specific band picked by CATSPERθ antibody. NC: none transfected. (*G*) Predicated 3D structure comparison of mouse CATSPERθ translated from ATG (171aa, magenta) and CTG (242aa, magenta, with extended N terminus in blue). (*H*) Fitting of predicated CTG initiated CATSPERθ 3D structure with the corresponding cryo-EM density map from different views.

### CTG start codon generates native full-length CATSPEPRθ in mouse testis

To better understand the sequence-structure-function relationship of CATSPERθ, we compared available CATSPERθ protein sequences from 162 different amniotes (**Fig. S2*A***). Markedly, the current NCBI annotated mouse CATSPERθ (171 residues, NP_001365176) is shorter than the homologs in most other species such as *Homo sapiens* (235 residues, NP_001239331.1) and *Ornithorhynchus anatinus* (platypus) (239 amino acids, XP_028918398) (**Fig. S2*A***). The genomic structure of *Catsperq* (*Tmem249*) is highly conserved, but the start site of the currently annotated mouse *Catsperq* in exon 3 is different from that of other species (**Fig. S2*B***). We considered the possibility that the difference in length of mouse CATSPERθ was due to incorrect annotation of its N terminus as it lacks a sequence portion conserved in multiple alignments (**Fig. S2*C***).

Indeed, by further investigating the 5’-UTR sequence of mouse *Catsperq*, a CTG was identified as a potential start codon. It is in the same reading frame as the annotated ATG-start codon (**Fig. 2*E***). The nucleotides at positions -3 and +4 of the CTG also match the preferred Kozak sequence. No frameshift mutations have been detected between the CTG and the downstream ATG among different rodent species that conserve both the CTG codon and the downstream sequences (**Fig. S2*D-E***). These lines of evidence suggest that the portion between CTG and ATG is a coding sequence. Compared to the downstream ATG-initiated CATSPERθ, the CTG-initiated mouse CATSPERθ has 242 amino acids by extending its N terminus with additional 71 amino acids. To validate whether mouse *Catsperq* uses this CTG as a start codon, constructs with different reading frames and sequence contexts were expressed in 293T cells (**Fig. 2*F***). The corresponding protein size was compared to that of native mouse CATSPERθ by Western blot probed with the CATSPERθ antibody, which specifically detects its C terminus (**Fig. 2*F***). The CTG codon in its native 5’-UTR context (plasmid CTG) was recognized as the start site as its protein product has the same apparent molecular weight as the positive control of plasmid CTG_R_ (CTG replaced by ATG). Most importantly, the translated protein has migrated to the same position on SDS-PAGE as the native mouse CATSPERθ, but not the ATG_1 or ATG_2 (a transcript variant annotated in the NCBI record) translated protein (**Fig. 2*F***). The use of the CTG start codon of *Catsperq* is likely shared with other species as their corresponding CATSPERθ shows high conservation (**Fig. S2*F and* S*3A and C***). The phylogenetic relations of this newly annotated full-length CATSPERθ are consistent with the organismal phylogeny (**Fig. S3*B***).

We compared the structure of the CTG-ORF and ATG-ORF encoded CATSPERθ predicted by AlphaFold (31) (**Fig. 2*G***). While both proteins contain two TM helixes, the first TM helix of the ATG initiated protein (15 residues) is shorter than the typical membrane-spanning helix (23 residues) and unconventionally includes the charged N terminus. However, the CTG initiated protein extends the first TM helix to 22 residues and the remaining 53 cytosolic residues interact with its own C-terminal cytosolic domain (**Fig. 2G**). The N-terminal extended CATSPERθ fits better to the protein density map corresponding to TMEM249 resolved by Lin et al. (19) (**Fig. 2*H***), compared to the currently annotated protein. Taken together, all these results strongly suggest that CATSPERθ is a CTG-initiated protein.

### CATSPERθ-deficient males are infertile due to defective sperm hyperactivation

To examine the functional significance of CATSPERθ in the CatSper complex, we created *Catsperq* knockout mice by CRISPR/Cas9 genome editing (**Fig. S4*A*-*D***). We obtained one mutant allele with two simultaneous deletions: 1,291bp deletion removing the region spanning the end of exon 1 to intron 5 plus 11bp deletion in exon 6, leading to an early terminational frameshift (**Fig. S4*B*-*C***). Western blot and immunocytochemistry analyses further validate the successful generation of *Catsperq* knockout mice by specific loss of CATSPERθ (**Fig. 3*A-B***). *Catsperq* knockout mice have no gross phenotype. There was no difference in epididymal sperm count or sperm morphology between *Catsperq*-null males and *wt* males (**Fig. S4*E*-*F***) or in histological examination of the testes (**Fig. S4*G***). *Catsperq*-null females have normal mating behavior and give birth to litters when mated with *wt* or *mono-allelic* heterozygous (*het*) males. However, *wt* or *het* females mated with *Catsperq*-null male mice produced no litters (**Fig. 3*C***, *left*). Consistently, *Catsperq*-null spermatozoa were unable to fertilize COC-intact oocytes *in vitro* (**Fig. 3*C***, *right*).

**Fig. 3.**
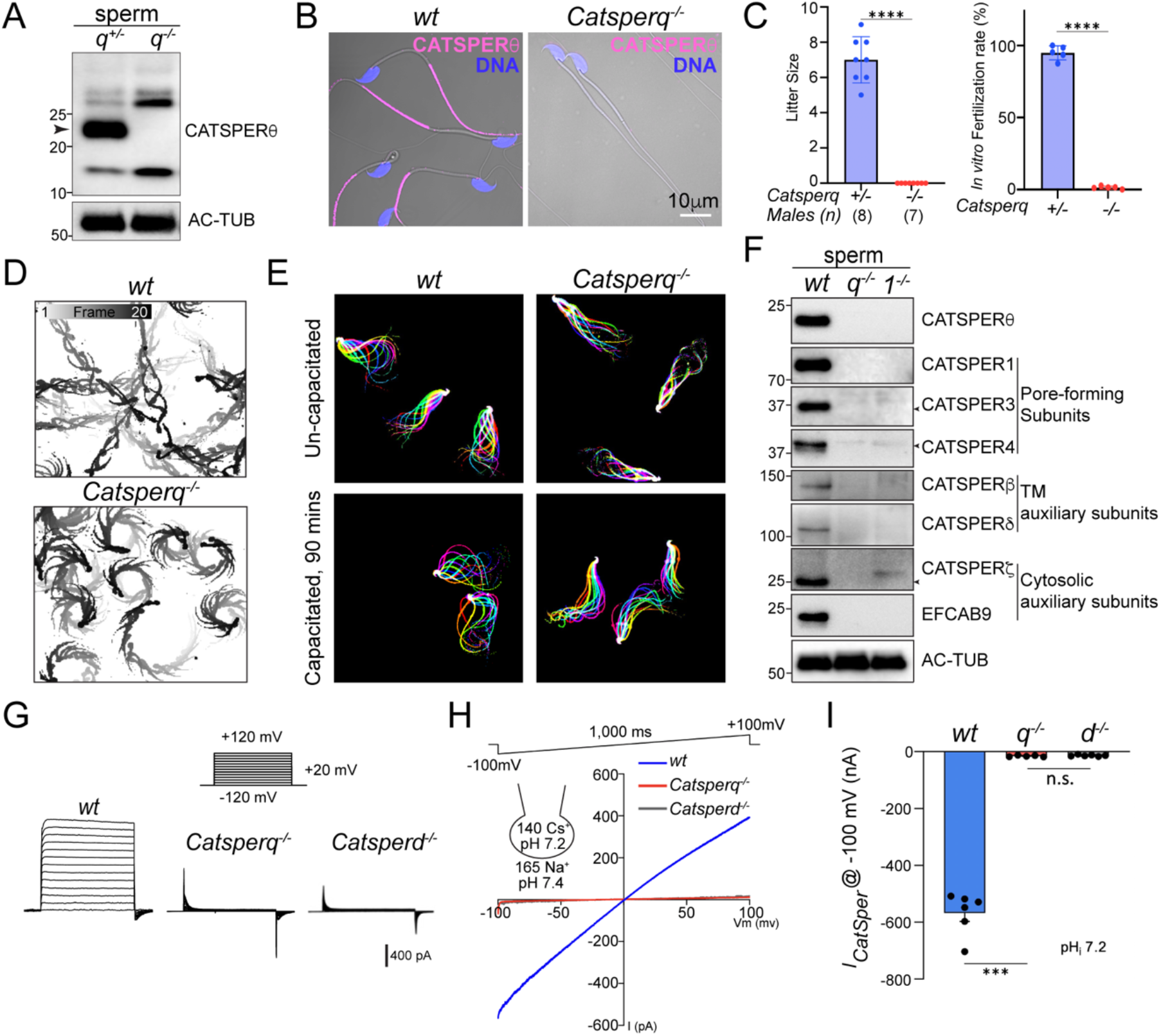
Genetic disruption of *Catsperq* renders males infertile due to defective sperm hyperactivation and the absence of *I*_*CatSper*_. (*A-B*) Validation of *Catsperq*^*-/-*^ mice generation by immunoblot analysis (*A*) and immunostaining (*B*) of CATSPERθ. Arrowhead in (*A*) indicates the corresponding protein band of CATSPERθ. (*C*) Fertility recording of natural mating and *in vitro* fertilization (IVF). 8 *Catsperq*^*+/-*^ and 7 *Catsperq*^*-/-*^ males were housed with *wt* females for 3 months, the litter size (*left*) and IVF rates of cumulus-oocyte complexes with *Catsperq*^*+/-*^ and *Catsperq*^*-/-*^ sperm (*right*) were recorded. ****P<0.0001. (*D*) Trajectory of free-swimming sperm from *wt* and *Catsperq*^*-/-*^ epididymal sperm after incubating in TYH medium for 10 min. Shown is time-scale bar(s). (*E*) Flagellar waveform analysis of uncapacitated (*upper*) and capacitated (*lower*) sperm from *wt* (*left*) and *Catsperq*^*-/-*^ (*right*) males. Movies were recorded at 200 fps from the head-tethered cells to glass coverslips before and after incubating under capacitation conidiations. (*F*) Absence of CatSper proteins in *Catsperq*^*-/-*^ sperm by immunoblot analysis. Arrows indicate the corresponding band of each protein. (*G*-*H*) Representative *I*_*CatSper*_ traces from *wt, Catsperq*^*-/-*^ and *Catsperd*^*-/-*^ sperm by step protocol (−120 to +120 mV) in 20 mV increments (*G*) and by voltage ramp (−100 to +100 mV) from 0 mV holding potential (*H*). The cartoon in (*H*) represents pH and monovalent ion composition in pipet and bath solutions. (*I*) Inward *I*_*CatSper*_ measured from *wt, Catsperq*^*-/-*^ (*q*^*-/-*^) and *Catsperd*^*-/-*^ (*d*^*-/-*^) sperm at -100mV. ***P<0.001. Data are presented as mean ± SEM (C and I).

We further examined sperm motility. We recorded free-swimming sperm trajectories after 10 min incubation under capacitating conditions (activated) and found that *wt* sperm swim relatively straight (**Fig. 3*D***, *upper*, **Movies S1**). By contrast, *Catsperq*-null sperm showed circulating swim path (**Fig. 3*D***, *lower*, **Movies S2**), a typical pattern observed for CatSper-deficient sperm that cannot roll along the axis (32) whereas the overall motility of *Catsperq*-null sperm is comparable to that of *wt* sperm (**Fig. S5*A***). Computer-assisted sperm analysis (CASA) further revealed that the mutant sperm have defects in sperm motility (**Fig. S5*B-E***). Specifically, total motility and progressive motility were dramatically reduced after incubating under capacitation conditions (**Fig. S5*A-B***), consistent with the previously reported impaired sperm motility maintenance in CatSper-deficient sperm (17, 24). The velocity of curvilinear (VCL), average path velocity (VAP), and straight-line velocity (VSL) were significantly reduced with *Catsperq*-null sperm than those of *wt* sperm, indicating defective hyperactivation of *Catsperq*-null sperm. Consistent with this idea, flagellar waveform analysis clearly demonstrated that capacitated *Catsperq*-null sperm failed to develop hyperactivation whereas *wt* sperm developed asymmetric beating with increased amplitude under the same conditions (**Fig. 3*E*, Movies S3-S6**). Taken together, our results show that the *Catsperq*-null male mice are infertile due to defective sperm hyperactivation, suggesting inactivation and/or absence of the CatSper channel.

### *Catsperq*-null sperm lack all the CatSper subunits and *I*_*CatSper*_

We first investigated whether the CatSper channel is present in *Catsperq*-null spermatozoa by examining the protein levels of the CatSper subunits. Immunoblot analyses clearly demonstrate that all the previously known CatSper components examined are missing in the absence of CATSPERθ, just as in *Catsper1*-null sperm (**Fig. 3*F***). Consistently, these CatSper subunits wereabsent or barely detectable by immunocytochemistry (**Fig. S5*F***). To test whether any small level of functional CatSper channel exists, we next performed whole sperm patch recording to measure *I*_*CatSper*_, the sperm-specific and pH-sensitive ion current mediated by the CatSper channel. *I*_*CatSper*_ is usually measured by Na^+^ conductance in divalent-free solution because it is small and difficult to record in normal bath solution containing 2 mM Ca^2+^ (33). We measured *I*_*CatSper*_ by keeping the intrapipette solution slightly alkaline at pH 7.2 to better visualize the small current if there is any. In *wt* spermatozoa, *I*_*CatSper*_ mediated a large inward current (−560 ± 73 pA at -100 mV). However, no current was detected in *Catsperq*-null spermatozoa, a similar level to that of *Catsperd*-null sperm (**Fig. 3*G-I***). Our results indicate that when *Catsperq* is inactivated, the sperm completely lacks functional CatSper channel due to the absence of CatSper proteins.

### CATSPERθ is required for CatSper channel assembly

Given that CATSPERθ is indispensable for functional CatSper channel in cauda epididymal sperm, we examined the total protein levels of each CatSper subunit in *Catsperq*-null testis. In the testis microsome, all the CatSper subunits examined have the same protein level as in *wt* (**Fig. 4*A***), suggesting that loss of CATSPERθ does not affect the protein expression and/or stability of other CatSper subunits in developing male germ cells. Nevertheless, CatSper components are unable to exit the cell body and be targeted to the spermatid flagella. As shown in **Fig. 4*B***, at step 10-12 of developing spermatid cells, we detected CATSPER1 in both the cell body and the prospective principal piece in *wt* spermatids, whereas the signal was only observed in the cell body in *Catsperq*-null spermatids. These results suggest that CATSPERθ is essential for the CatSper complex assembly and/or trafficking in spermatids.

**Fig. 4.**
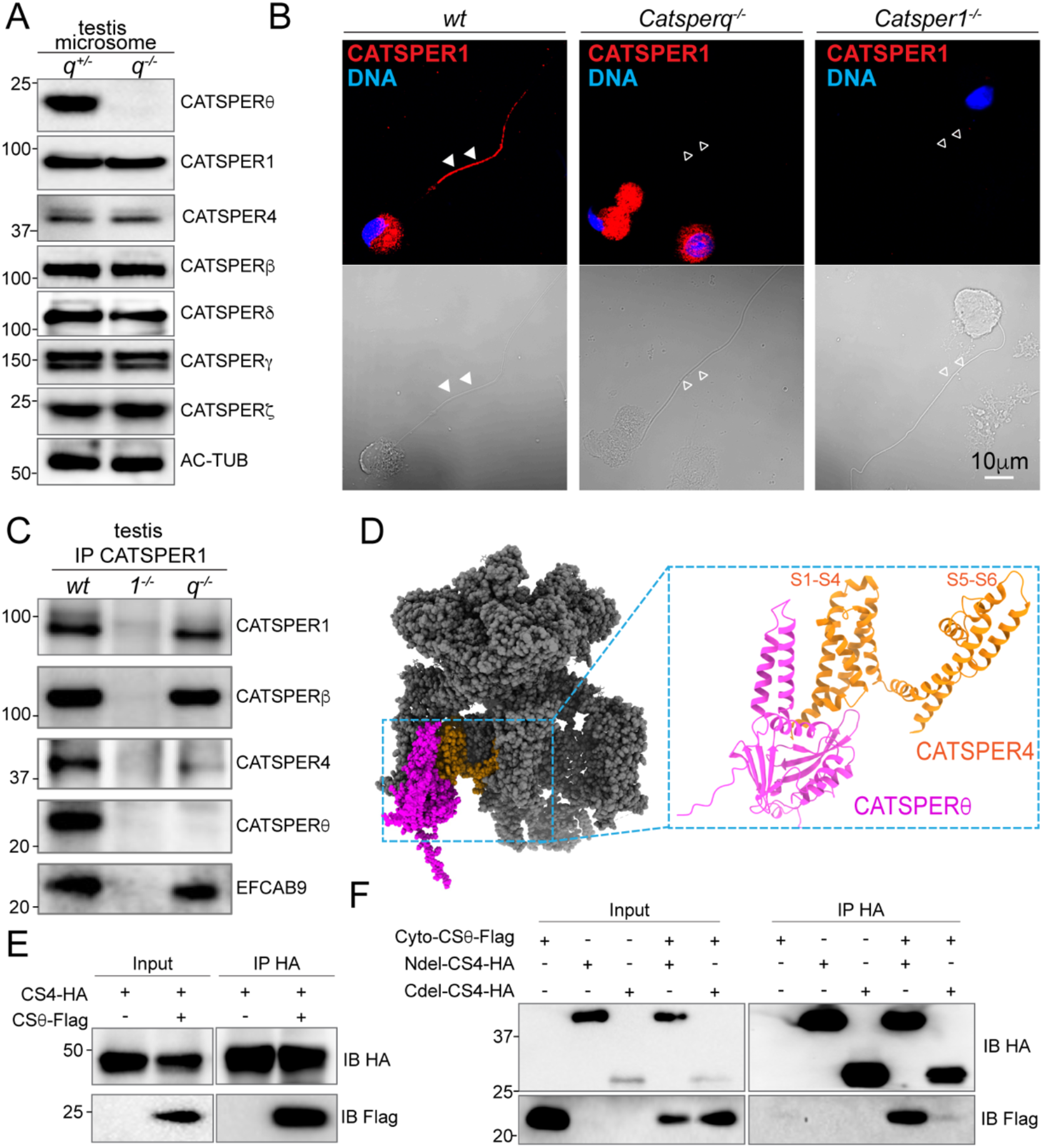
CATSPERθ loss-of-function compromises CATSPER4 assembly into CatSper Complex. (*A*) Immunoblot analysis of CatSper subunits in *Catsperq*^*-/-*^ and *wt* testis microsome. (*B*) Immunostaining of CATSPER1 in developing male germ cells at various stages isolated from testes of *wt* (*left*), *Catsperq*^*-/-*^ (*middle*), and *Catsper1*^*-/-*^ (*right*) males. The corresponding DIC images were shown (*lower*). Filled arrowheads indicate the presence of signal in the tail; empty arrowheads indicate the absence of the signal. (*C*) Proteins solubilized from *wt, Catsper1*^*-/-*^ and *Catsperq*^*-/-*^ testes microsome were immunoprecipitated (IP) with anti-CATSPER1 antibody and followed with immunoblot analysis with indicated antibodies. (*D*) The structure of mouse CatSper complex is modified from (19) in sphere presentation, with CATSPERθ in magenta, CATSPER4 in yellow and all other subunits in grey. Of note is that the structure of CATSPER4 here are mainly the TM domains (S1-S6), due to the N and C cytosolic domains of it are not resolved in the CatSper automatic structure. Zoomed CATSPERθ and CATSPER4 are in ribbon presentation. (*E*) CATSPER4-HA (CS4-HA) and/or CATSPERθ-Flag (CSθ-Flag) were transfected into 293T cells and precipitated with anti-HA antibodies. The input and immunocomplexes were blotted with anti-HA and anti-Flag antibodies. (*F*) two TM domain deleted CATSPERθ (Cyto-CSθ-Flag) and or truncated CATSPER4 (N-terminal cytosolic domain deleted: Ndel-CS4-HA; C-terminal cytosolic domain deleted: Cdel-CS4-HA) were transfected into 293T cells and precipitated with anti-HA antibodies. The input and immunocomplexes were blotted with anti-HA and anti-Flag antibodies.

Next, we examined the assembly of the CatSper channel in the testis. Co-immunoprecipitation (co-IP) with anti-CATSPER1 antibody showed that, unlike all the other CatSper subunits examined, the protein level of CATSPER4 in the CatSper complex is significantly compromised in the absence of CATSPERθ (**Fig. 4*C***), suggesting that CATSPERθ functions to stabilize CATSPER4. The proximity between CATSPER4 and CATSPERθ in the cryo-EM structure of CatSper also supports this hypothesis (**Fig. 4*D***). Therefore, we investigated the interaction between CATSPERθ and CATSPER4 in heterologous 293T cells by co-IP experiments. When co-expressed, full length CATSPERθ is indeed complexed with full length CATSPER4 (**Fig. 4*E***). Based on the cryo-EM structure of CatSper, it is likely that the interaction is mainly mediated by the cytosolic domain of each protein. To test this idea and further narrow down which part of CATSPER4 mediates this interaction, co-IP was performed with the cytosolic domain of CATSPERθ (Cyto-CSθ), N-terminal cytosolic domain deleted CATSPER4 (Ndel-CS4) and C-terminal cytosolic domain deleted CATSPER4 (Cdel-CS4) (**Fig. 4*F***). The interaction between Ndel-CS4 and Cyto-CSθ is maintained, but the CATSPER4 interaction with Cyto-CSθ is largely disrupted when its C-terminal cytosolic domain is removed. These results indicate that the CATSPERθ-CATSPER4 interaction is mainly mediated by the cytosolic domain of CATSPERθ and the C-terminal cytosolic domain of CATSPER4. The remaining weak interaction observed between Ndel-CS4 and Cyto-CSθ suggests that the intracellular loops between the TM domains of CATSPER4 may also be involved in the interaction between them. These results suggest that CATSPERθ efficiently stabilizes CATSPER4 assembled into the CatSper complex by serving a scaffold for CATSPER4.

### CATSPERθ is capable of self-interaction to form CatSper dimer

Notably, the assembly of CATSPER4 into the CatSper complex is not completely abolished in *Catsperq*-null spermatids (**Fig. 4*C***). However, the remaining CATSPER4-containing CatSper complex is still unable to leave the cell body to reach the sperm flagella (**Fig. 4*B*** and **Fig. 3**), indicating that the defects in *Catsperq*-null spermatids are beyond the level of monomeric CatSper complex assembly. Our recent application of cryo-ET to intact mouse and human spermatozoa revealed the 3D structure and higher-order organization of the native CatSper channels. The repeated CatSper units are interconnected and form zigzag rows on the extracellular side (**Fig. S6*A***, *left*). Underneath the zigzag rows, the CatSper form diagonal arrays on the intracellular side (26) (**Fig. S6*A*** *right*). Each diagonal stripe is comprised of two connected intracellular structures from two staggered CatSper channel units, suggesting that CatSper dimers (each diagonal stripe) are the building blocks for the zigzag rows (26). Together with CATSPERη (TMEM262), another newly annotated three TM domain containing CatSper component revealed by Lin et al. (19), CATSPERθ is also located at the interface within the CatSper dimer in which CatSper4/β side position face-to-face with 180° rotation (26) (**Fig. 5*A*** and **Fig. S6*A***). Thus, we hypothesize that CATSPERθ plays a role in CatSper dimer formation by connecting two complex units. Without CATSPERθ, the CatSper dimer is unlikely to form properly, preventing the channel from trafficking to the flagellar membrane and forming the higher-order arrangement. We tested this idea by examining whether CATSPERθ can self-interact. We expressed recombinant CATSPERθ with either HA or Flag tag in 293T cells and found that they form immunocomplex with each other (**Fig. 5*B***), supporting its contribution to CatSper dimer formation.

**Fig. 5.**
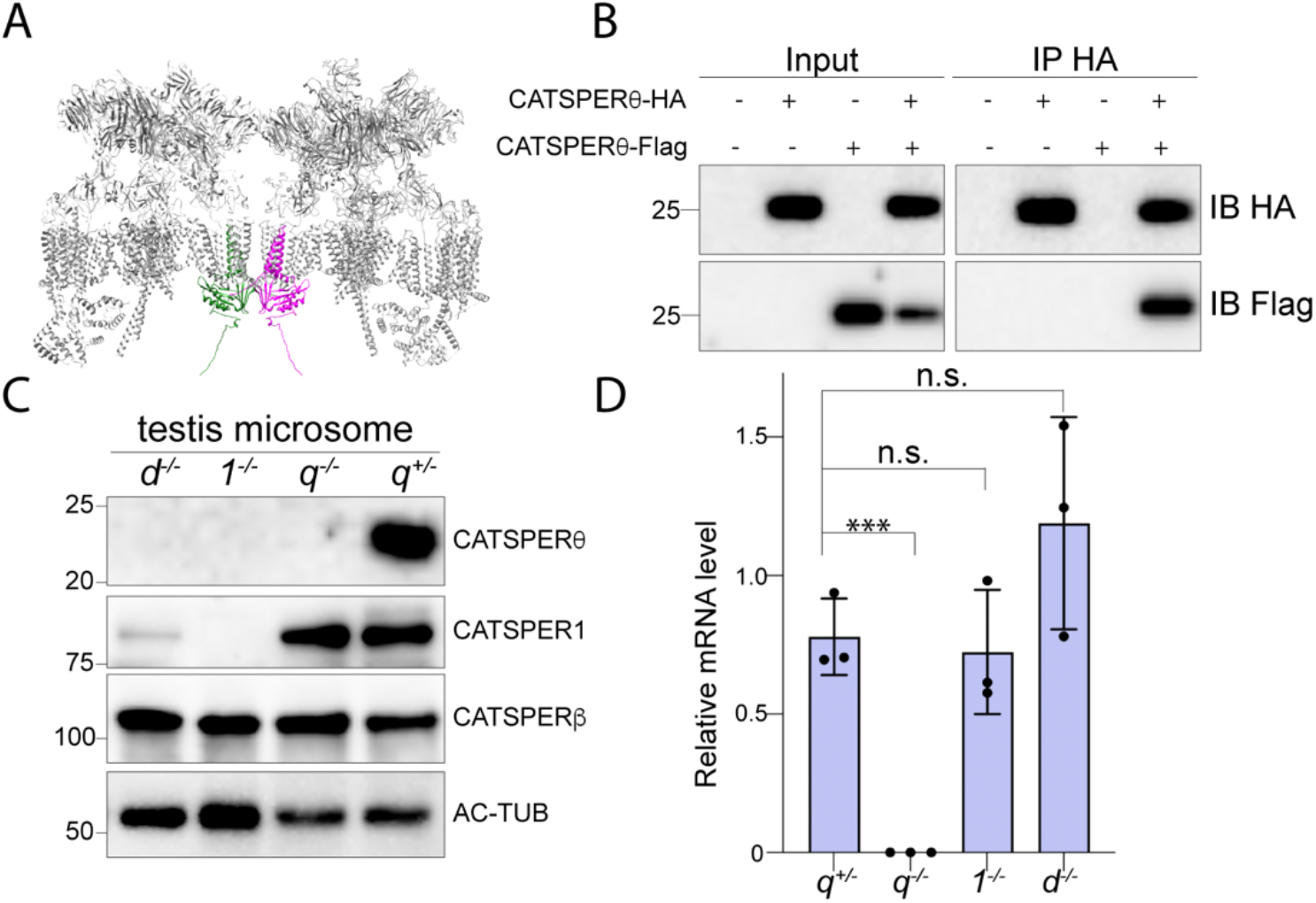
CATSPERθ participated in CatSper dimer formation and works as a checkpoint. (*A*) Side view of the pseudo-atomic model of a CatSper complex dimer showing potential interactions between two CATSPERθ. Two CATSPERθ from two CatSper units are highlighted with green and magenta respectively. (*B*) CATSPERθ-HA and/or CATSPERθ-Flag were transfected into 293T cells and precipitated with anti-HA antibodies. The immunocomplexes were blotted with anti-HA and anti-Flag antibodies. (*C*) Immunoblot analysis of indicated CatSper subunits in the solubilized testis microsome from various CatSper TM subunits knockout mice. (*D*) *Catsperq* mRNA expression level in mice testes is measured by real time RT-PCR. TATA binding protein (TBP) is used as a control for normalization. ***P<0.001. Data are presented as mean ± SEM, n=3.

### CatSper TM subunit deficiency results in loss of CATSPERθ during spermatogenesis

Genetic abrogation of any of the CatSper pore-forming (CATSPER1-4) or auxiliary TM subunits (i.e., CATSPERθ) results in loss of the entire CatSper channel complex in mature mouse spermatozoa (17, 23, 27, 34). These results indicate that CatSper complex missing any of these essential TM subunits are unable to traffic to the sperm flagellum during tail formation. This phenomenon implies the existence of quality control checkpoint for properly assembled CatSper complex and those without complete CatSper TM subunits are retained in the cell body. Loss of CATSPERθ leads to the same consequence (**Fig. 3**). The spermatids lacking the TM subunits of the CatSper channel may have common defects. To test this hypothesis, we evaluated the protein levels of different TM subunits in *Catsper1*-null, *d*-null, or *q*-null testes compared to those in *Catsperq*-het testes. The testicular CATSPER1 level is not affected by abrogation of *Catsperq*, distinct from the compromised level in *Catsperd*-null males (23). The amount of CATSPERγ also remains unchanged in the testes of all the genotypes tested. Intriguingly, CATSPERθ is not detected in the testes of *Catsperd*-null or *Catsper1*-null males (**Fig. 5*C*** and **Fig. S6*B***). Since *Catsperq* mRNA levels in *Catsper1*-null and *d*-null testes are not significantly different from that in *Catsperq*-het testis (**Fig. 5*D***), the absence of other TM subunits is likely to affect the *Catsperq* translation or the protein stability. Thus, the CATSPERθ levels may serve as a quality control checkpoint for the CatSper complex assembly. Only the complete CatSper complex with all TM subunits would associate with CATSPERθ to form a dimer and traffics to the flagella.

### CATSPERθ is actively processed during sperm capacitation

To explore a physiological function of CATSPERθ in the zigzag channel dimers in the regulation of sperm motility, we next examined the levels of CATSPERθ before and after inducing capacitation. As previously reported, CATSPER1 is processed during sperm capacitation, whereas other subunits including CATSPERγ shows no obvious changes in the protein level (27, 30) (**Fig. S7*A-C***). Interestingly, like CATSPER1, CATSPERθ is significantly decreased after incubating for 90 min under capacitating conditions. To test whether this change reflects sperm motility status, we performed motility correlated imaging of individual sperm cells (27). In vitro capacitated spermatozoa placed on photo-etched grid coverslips were videotaped to record their motility patterns before using the same coverslips to locate spermatozoa and correlate CASTSPERθ levels with the motility patterns (**Fig. S7*D***). We found that spermatozoa that were motile (either hyperactivated or vibrating) retained an intact CATSPERθ signal. In contrast, CATSPERθ was not detected in a significant proportion of immotile spermatozoa. These results suggest that sperm motility is impaired when the CatSper dimers are disrupted by CATSPERθ processing.

## Discussion

### The non-canonical CTG start codon of CATSPERθ

It has long been thought that eukaryotic translation almost always initiates at an AUG start codon. However, recent advances in ribosome footprint mapping have revealed that non-AUG start codons (e.g., CUG, GUG, and UUG) are also used at an unexpectedly high frequency, with CUG generally being the most efficient among the non-AUG start codons (35). The use of CUG has been found to generate protein isoforms. For example, thioredoxin-glutathione reductase 3 (TXNRD3), a member of the thioredoxin reductase family that is particularly abundant in elongating spermatids (36), uses the CUG start codon to generate the long isoform in mouse testis (37). The oncogenic transcription factor MYC also uses the CUG codon to generate an N-terminally extended isoform of c-Myc (38), just as the use of the CUG as start codon extends the N-terminus of CATSPERθ with respect to the use of the downstream AUG codon. The fit of this N-terminally extended CATSPERθ with the cryo-EM density map supports the use of CUG as the start codon for mouse *Catsperq*. However, unlike the above-mentioned TXNRD3 or MYC, the use of the CUG start codon does not generate another *Catsperq* isoform. This possibility is ruled out by the fact that the antibody raised against the very C-terminal region of CATSPERθ only recognize a single protein band.

Currently established models of translation suggest that initiation at non-AUG start codons is mediated by the methionine charged initiator tRNA (Met-tRNA_i_^Met^) through ‘wobble’ interactions with the anticodon (39). However, Leucine-tRNA was also found to initiate at CUG start codons for synthesizing and presenting new proteins by major histocompatibility complex in the immune response (40). Which initiator tRNA, Met-tRNA or Leu-tRNA, is used to decode the CUG start codon of *Catsperq* in testis remains to be further investigated.

### CATSPERθ functions in assembling monomeric CatSper complex

The cryo-EM structure of the isolated mouse CatSper complex visualized that all the previously known Type I single TM CatSper auxiliary subunits with large extracellular domain bind to each of the pore-forming subunits in a specific configuration: CATSPERβ, γ, δ and ε form a pavilion-like structure by pairing with each of the pore subunits CATSPER4, 1, 3 and 2, respectively (19). CATSPERθ stands out among the TM auxiliary subunits in that it contains two TM domains and a cytosolic globular domain. We propose that one role of CATSPERθ is to stabilize CATSPER4 in a single CatSper complex unit via its cytosolic regions. Notably, CATSPERη, another TM auxiliary subunit with three TM domains, is also located in close proximity to CATSPERβ that pairs with CATSPER4. Together with the previously suggested potential TM interaction between CATSPERθ and CATSPERη (19), the stability of CATSPER4 within a monomeric CatSper complex unit is likely also supported by CATSPERη and CATSPERβ. The specific localization of CATSPERθ at the center of a CatSper dimer and its self-interaction ability make it a good candidate to be responsible for CatSper dimer formation. Yet, there are additional components that are likely to be involved in the dimer formation including the extracellular CATSPERβ-CATSPERβ interaction at the intradimer interface and the unresolved protein density identified in the center of the diagonal array (26), which need to be further investigated.

### CATSPERθ serves as a checkpoint for CatSper complex assembly

The cytosolic auxiliary subunits (CATSPERζ, EFCAB9 or CATSPERτ) of the CatSper channel complex function mainly in modulating channel density or (in)activation (41). In the absence of either one of the cytosolic auxiliary subunits, the CatSper channel still assembles properly, targets to the flagellar membrane, and conducts *I*_*CatSper*_ albeit with reduced current density and/or altered sensitivity to intracellular pH and Ca^2+^ (22, 24, 25). In contrast, not only the pore-forming CATSPER1-4 but also the TM ancillary subunits are essential for the presence of the functional channel in the sperm flagella (23). The present study further supports this general notion by functionally annotating a novel CatSper component CATSPERθ into such an essential protein group. Disruption of *Catsperq* renders the entire CatSper complex completely missing in the epididymal sperm flagella, indicating its fundamental role in the CatSper channel assembly and/or trafficking.

We found that the absence of CATSPERθ during spermatogenesis is a common denominator for the mouse models that lack either a pore-forming or an ancillary TM subunit of the CatSper channel. We propose that CATSPERθ is a checkpoint to quality control CatSper assembly. In this regard, it is intriguing that *Catsperq* uses the noncanonical CTG start codon, which might be tightly regulated by the upstream CatSper assembly event. While the detailed mechanisms require further investigation, the presence of CATSPERθ is likely to be a key quality control checkpoint that signals “go” for the properly assembled CatSper complex to traffic to the flagellum (**Fig. S7*E***). Additionally, our results show that capacitation-induced processing of CATSPERθ are likely to affect the higher-order arrangement of CatSper and the channel activity.

In summary, using coevolutionary and biochemical analyses combined with mouse genetics, we have structurally and functionally annotated a CUG initiated *Tmem249*-encoded CATSPERθ as an essential CatSper component that functions in CatSper complex assembly. CATSPERθ serves as a scaffold platform to stabilize CATSPER4 and contributes to linking two monomeric CatSper complexes into a dimer. Loss of CATSPERθ prevents the assembled CatSper complex from exiting the cell body and renders male mice infertile, suggesting a fundamental role for CATSPERθ as a checkpoint for the higher-order assembly.

## Materials and Methods

### Identification of CatSper components by coevolutionary analysis

We performed a global analysis of coevolutionary associations among 60675 orthologous gene groups from 1256 eukaryotic genomes as classified by OrthoDB (Ver. 10). Coevolution among different orthogroups was assessed through a procedure especially suited to reveal associations between genes with a discontinuous pattern of presence/absence as observed in CatSper (42, 43), we enumerated concordant transitions (1→0 or 0→1) in pairs of binary vectors describing the presence/absence of orthologous genes in complete genomes ordered according to species phylogeny, so that concordant state transitions correspond to simultaneous events of gene loss or gain. Significant pairwise associations (adjusted P-value < 0.001; Fisher exact test with Bonferroni correction) were clustered with MCL (Markov cluster) (44) and the resulting clusters were parsed for the presence of known CatSper components. Nine CatSper components (*Catsper* 1-4, *b, g, d/e, and Efcab9*) were found in a single MCL cluster which additionally contained the uncharacterized gene *Tmem249*. The procedure was carried out using Python and R scripts on a 64-core HPC node with 384 Gb RAM.

### Generation of *Catsperq*^*-/-*^ mice by CRISPR/Cas9 and genotyping

The *Catsperq*^*-/-*^ mice were generated using CRISPR/Cas9 based gene targeting. Two guide RNAs (gRNAs) gRNA1 (5’-accctggaattgtctcgggt-3’) and gRNA2 (5’-taagtcaagtgtcaacgacc-3’) targeting the coding region of *Catsperq* were designed using CRISPR direct software (http://crispr.dbcls.jp/) (45). Zygotes were isolated on the day of the coagulation plug (= E0.5) from super ovulated female mice (B6D2F1) mated with B6D2F1 males. To remove cumulus cells, zygotes were incubated in hyaluronidase solution (0.33 mg/mL) (Sigma-Aldrich). Ribonucleoprotein complexes containing synthesized CRISPR RNA (crRNA) (Sigma-Aldrich), trans-activating crRNA (tracrRNA) (TRACRRNA05N-5NMOL, Sigma-Aldrich), and CAS9 protein (A36497, Thermo Fisher Scientific) were electroporated into fertilized eggs using a NEPA21 super electroporator (NEPA GENE) (46). Electroporated zygotes were incubated in potassium simplex optimization medium (KSOM) (47) at 37°C and 5% CO_2_ until the next day. Two-cell embryos were transferred into the oviducts of pseudo pregnant recipient females. The resulting pups were genotyped using primers TMEM249_F (5’-tgtggtcaatagaaaagcccct-3’), TMEM249_R (5’-cgcgtctcctcccacaagtac-3’) and TMEM249_R’ (5’-aaaggaggccagggctcaggcccca-3’).

### Preparation and solubilization of testis microsome and immunoprecipitation

Mouse testes were homogenized in 0.32M sucrose and cell debris and nuclei were removed by centrifugation at 1,000 x g, 4°C for 20 min. Supernatant was centrifuged at 100,000 x g, 4°C for 60 min to collect membrane microsome. The microsome fraction was solubilized in 1% Triton X-100 in PBS (10 mg microsome per ml) with protease inhibitor cocktail (complete mini, Roche) by rocking at 4°C for 60 min. The lysate was cleared by centrifugation at 18,000 x g, 4°C for 30 min to obtain solubilized proteins from the supernatant. The supernatant was immunoprecipitated with anti-CATSPER1 antibody and Protein A magnetic beads (BIO-RAD) with the same method for recombinant protein co-IP. Proteins were detected with corresponding rabbit raised primary antibodies and horseradish peroxidase-conjugated goat-anti-rabbit secondary antibody.

### Sperm immunocytochemistry

Sperm cells from the cauda epididymis attached to glass coverslips were fixed in 4% paraformaldehyde in PBS for 20 min, permeabilized with 0.1% Triton X-100 in PBS for 10 mins, washed with PBS and blocked with 10% goat serum for 1h. Samples were stained overnight with indicated primary antibody in 10% goat serum at 4°C. For secondary antibodies, Alexa568 or Alexa488 conjugated goat-anti-rabbit (Invitrogen) were used. Images were acquired by confocal microscopy (Zeiss LSM710 Elyra P1).

### Preparation of spermatogenic cells for immunocytochemistry

Testes from adult male mice were collected and tunica albuginea was removed to harvest seminiferous tubules. Collected seminiferous tubules were washed with ice-cold PBS two times and chopped into small pieces to release germ cells. Chopped mixture was filtered with 40 mm Nylon Mesh cell strainer to get dissociated germ cells. Filtered testicular cells were used for immunocytochemistry.

### Mating test and in vitro fertilization

Sexually matured male mice were individually caged with three 8-wk-old B6D2F1 female mice for 2 months. Plugs were checked every morning and the number of pups was counted on the day of birth. For in vitro fertilization, spermatozoa were collected from the cauda epididymis and incubated in a TYH drop (48) for 120 min at 37°C under 5% CO_2_. Eggs were collected from super ovulated females. The spermatozoa were then added to the TYH drop containing intact eggs at a final concentration of 2×10^5^ spermatozoa/ml. Two-cell embryos were counted the next day.

### Flagellar waveform analysis

To tether sperm head for planar beating, non-capacitated or capacitated spermatozoa (2×10^5^ cells) from adult male mice were transferred to the fibronectin-coated 37°C chamber for Delta T culture dish controller (Bioptechs) filled with HEPES-buffered HTF medium (H-HTF) for 1 min. Flagellar movements of the tethered sperm were recorded for 2 s with 200 fps using pco.edge sCMOS camera equipped in Axio observer Z1 microscope (Carl Zeiss). All movies were taken at 37°C within 10 min after transferring sperm to the imaging dish. FIJI software was used to measure beating frequency of sperm tail, and to generate overlaid images to trace waveform of sperm flagella as previously described (22).

### Sperm free swimming tracing

Cauda epididymal spermatozoa were dispersed in a drop of TYH medium and incubated for 10 min and 2 hr at 37°C under 5% CO_2_. Spermatozoa were collected from the top of the drop and observed with an Olympus BX-53 microscope equipped with a high-speed camera (HAS-L1, Ditect).

### Electrophysiology

Corpus epididymal sperm were washed and resuspended in HS medium followed by attached on a 35mm culture dish. A Gigaohm seal was formed at the cytoplasmic droplet of sperm (33). The cell was broken in by applying voltage pulses (450-600 mV, 5ms) and simultaneous suction. Whole-cell CatSper currents were recorded from *wt, CatSperd*^*-/-*^, *and CatSperq*^*-/-*^ sperm in divalent-free bath solution (DVF, 150mM Na gluconate, 20mM HEPES, and 5mM Na_3_HEDTA, pH 7.4). Intrapipette solution for CatSper current recording consists of 135mM CsMes, 10mM HEPES, 10mM EGTA, and 5mM CsCl adjusted to pH 7.2 with CsOH. Data were sampled at 10 kHz and filtered at 2 kHz. The current data were analyzed with Clampfit (Axon, Gilze, Netherlands), and figures were plotted with Grapher 8 software (Golden Software, Inc., Golden, Colorado).

### Quantification and statistical analysis

Statistical analyses were performed using Student’s t test or two-way analysis of variance (ANOVA). Differences were considered significant at *p < 0.05, **p < 0.01, ***p < 0.001 and ****p < 0.0001.

## Acknowledgments

We thank Dr. Yanhe Zhao for this guidance in the fitting of CatSper cryo-EM structure to the cryo-ET structure and Ms. Eri Hosoyamada for technical assistance. This work was supported by start-up funds from National Institute of Child Health and Human Development (R01HD096745) and Grantham Foundation to J.-J.C.; the Ministry of Education, Culture, Sports, Science, and Technology (MEXT)/Japan Society for the Promotion of Science (JSPS) KAKENHI grants (JP22H03214 to H.M., JP19H05750 and JP21H05033 to M.I.) and National Institute of Child Health and Human Development (R01HD088412 and P01HD087157) to M.I.; the Italian Ministry for Education, University and Research (MIUR) PRIN grant 2017483NH8 to R.P.; the computational screening benefited from the HPC@Unipr and COMP-R frameworks; X.H. is a recipient of postdoctoral fellowships from Yale University School of Medicine James Hudson Brown-Alexander Brown Coxe Postdoctoral Fellowships in the Medical Sciences and The Lalor Foundation Fellowship.

## References

1. H. C. Lai, L. Y. Jan, The distribution and targeting of neuronal voltage-gated ion channels. Nat Rev Neurosci 7, 548–562 (2006).

2. A. M. Katz, Cardiac ion channels. N Engl J Med 328, 1244–1251 (1993).

3. E. A. Pereira da Silva, M. Martin-Aragon Baudel, M. F. Navedo, M. Nieves-Cintron, Ion channel molecular complexes in vascular smooth muscle. Front Physiol 13, 999369 (2022).

4. J. J. Devaux, K. A. Kleopa, E. C. Cooper, S. S. Scherer, KCNQ2 is a nodal K+ channel. J Neurosci 24, 1236–1244 (2004).

5. T. Tanabe et al., Primary structure of the receptor for calcium channel blockers from skeletal muscle. Nature 328, 313–318 (1987).

6. M. Takahashi, M. J. Seagar, J. F. Jones, B. F. Reber, W. A. Catterall, Subunit structure of dihydropyridine-sensitive calcium channels from skeletal muscle. Proc Natl Acad Sci U S A 84, 5478–5482 (1987).

7. G. W. Abbott, Kv Channel Ancillary Subunits: Where Do We Go from Here? Physiology (Bethesda) 37, 0 (2022).

8. Y. Kise et al., Structural basis of gating modulation of Kv4 channel complexes. Nature 599, 158–164 (2021).

9. K. Li et al., Tetrameric Assembly of K(+) Channels Requires ER-Located Chaperone Proteins. Mol Cell 65, 52–65 (2017).

10. A. Kosolapov, L. Tu, J. Wang, C. Deutsch, Structure acquisition of the T1 domain of Kv1.3 during biogenesis. Neuron 44, 295–307 (2004).

11. J. M. Robinson, C. Deutsch, Coupled tertiary folding and oligomerization of the T1 domain of Kv channels. Neuron 45, 223–232 (2005).

12. C. L. Anderson et al., Most LQT2 mutations reduce Kv11.1 (hERG) current by a class 2 (trafficking-deficient) mechanism. Circulation 113, 365–373 (2006).

13. C. L. Anderson et al., Large-scale mutational analysis of Kv11.1 reveals molecular insights into type 2 long QT syndrome. Nat Commun 5, 5535 (2014).

14. G. L. Lukacs, A. S. Verkman, CFTR: folding, misfolding and correcting the DeltaF508 conformational defect. Trends Mol Med 18, 81–91 (2012).

15. S. S. Suarez, S. M. Varosi, X. Dai, Intracellular calcium increases with hyperactivation in intact, moving hamster sperm and oscillates with the flagellar beat cycle. Proc Natl Acad Sci U S A 90, 4660–4664 (1993).

16. D. Ren et al., A sperm ion channel required for sperm motility and male fertility. Nature 413, 603–609 (2001).

17. H. Qi et al., All four CatSper ion channel proteins are required for male fertility and sperm cell hyperactivated motility. Proc Natl Acad Sci U S A 104, 1219–1223 (2007).

18. T. A. Quill, D. Ren, D. E. Clapham, D. L. Garbers, A voltage-gated ion channel expressed specifically in spermatozoa. Proc Natl Acad Sci U S A 98, 12527–12531 (2001).

19. S. Lin, M. Ke, Y. Zhang, Z. Yan, J. Wu, Structure of a mammalian sperm cation channel complex. Nature 595, 746–750 (2021).

20. J. Liu, J. Xia, K. H. Cho, D. E. Clapham, D. Ren, CatSperbeta, a novel transmembrane protein in the CatSper channel complex. J Biol Chem 282, 18945–18952 (2007).

21. H. Wang, J. Liu, K. H. Cho, D. Ren, A novel, single, transmembrane protein CATSPERG is associated with CATSPER1 channel protein. Biol Reprod 81, 539–544 (2009).

22. J. J. Chung et al., CatSperzeta regulates the structural continuity of sperm Ca2+ signaling domains and is required for normal fertility. Elife 6 (2017).

23. J. J. Chung, B. Navarro, G. Krapivinsky, L. Krapivinsky, D. E. Clapham, A novel gene required for male fertility and functional CATSPER channel formation in spermatozoa. Nat Commun 2, 153 (2011).

24. J. Y. Hwang et al., Dual Sensing of Physiologic pH and Calcium by EFCAB9 Regulates Sperm Motility. Cell 177, 1480–1494 e1419 (2019).

25. J. Y. Hwang, H. Wang, Y. Lu, M. Ikawa, J. J. Chung, C2cd6-encoded CatSpertau targets sperm calcium channel to Ca(2+) signaling domains in the flagellar membrane. Cell Rep 38, 110226 (2022).

26. Y. Zhao et al., 3D structure and in situ arrangements of CatSper channel in the sperm flagellum. Nat Commun 13, 3439 (2022).

27. J. J. Chung et al., Structurally distinct Ca(2+) signaling domains of sperm flagella orchestrate tyrosine phosphorylation and motility. Cell 157, 808–822 (2014).

28. T. A. Quill et al., Hyperactivated sperm motility driven by CatSper2 is required for fertilization. P Natl Acad Sci USA 100, 14869–14874 (2003).

29. C. Wiesehofer et al., CatSper and its CaM-like Ca(2+) sensor EFCAB9 are necessary for the path chirality of sperm. FASEB J 36, e22288 (2022).

30. L. Ded, J. Y. Hwang, K. Miki, H. F. Shi, J. J. Chung, 3D in situ imaging of the female reproductive tract reveals molecular signatures of fertilizing spermatozoa in mice. Elife 9 (2020).

31. J. Jumper et al., Highly accurate protein structure prediction with AlphaFold. Nature 596, 583–589 (2021).

32. K. Miki, D. E. Clapham, Rheotaxis guides mammalian sperm. Curr Biol 23, 443–452 (2013).

33. Y. Kirichok, B. Navarro, D. E. Clapham, Whole-cell patch-clamp measurements of spermatozoa reveal an alkaline-activated Ca2+ channel. Nature 439, 737–740 (2006).

34. H. Wang, L. L. McGoldrick, J. J. Chung, Sperm ion channels and transporters in male fertility and infertility. Nat Rev Urol 18, 46–66 (2021).

35. M. G. Kearse, J. E. Wilusz, Non-AUG translation: a new start for protein synthesis in eukaryotes. Genes Dev 31, 1717–1731 (2017).

36. H. Wang et al., Redox regulation by TXNRD3 during epididymal maturation underlies capacitation-associated mitochondrial activity and sperm motility in mice. J Biol Chem 298, 102077 (2022).

37. M. V. Gerashchenko, D. Su, V. N. Gladyshev, CUG start codon generates thioredoxin/glutathione reductase isoforms in mouse testes. J Biol Chem 285, 4595–4602 (2010).

38. S. R. Hann, M. W. King, D. L. Bentley, C. W. Anderson, R. N. Eisenman, A non-AUG translational initiation in c-myc exon 1 generates an N-terminally distinct protein whose synthesis is disrupted in Burkitt’s lymphomas. Cell 52, 185–195 (1988).

39. D. S. Peabody, Translation Initiation at Non-Aug Triplets in Mammalian-Cells. Journal of Biological Chemistry 264, 5031–5035 (1989).

40. S. R. Starck et al., Leucine-tRNA Initiates at CUG Start Codons for Protein Synthesis and Presentation by MHC Class I. Science 336, 1719–1723 (2012).

41. J. Y. Hwang, J. J. Chung, CATSPER Calcium Channels: twenty years on. Physiology (Bethesda) 10.1152/physiol.00028.2022 (2022).

42. G. Dey, A. Jaimovich, S. R. Collins, A. Seki, T. Meyer, Systematic Discovery of Human Gene Function and Principles of Modular Organization through Phylogenetic Profiling. Cell Rep 10, 993–1006 (2015).

43. S. Cokus, S. Mizutani, M. Pellegrini, An improved method for identifying functionally linked proteins using phylogenetic profiles. BMC Bioinformatics 8 Suppl 4, S7 (2007).

44. A. J. Enright, S. Van Dongen, C. A. Ouzounis, An efficient algorithm for large-scale detection of protein families. Nucleic Acids Res 30, 1575–1584 (2002).

45. Y. Naito, K. Hino, H. Bono, K. Ui-Tei, CRISPRdirect: software for designing CRISPR/Cas guide RNA with reduced off-target sites. Bioinformatics 31, 1120–1123 (2015).

46. F. Abbasi et al., RSPH6A is required for sperm flagellum formation and male fertility in mice. J Cell Sci 131 (2018).

47. Y. Ho, K. Wigglesworth, J. J. Eppig, R. M. Schultz, Preimplantation development of mouse embryos in KSOM: augmentation by amino acids and analysis of gene expression. Mol Reprod Dev 41, 232–238 (1995).

48. Y. Muro et al., Behavior of Mouse Spermatozoa in the Female Reproductive Tract from Soon after Mating to the Beginning of Fertilization. Biol Reprod 94, 80 (2016).

