## Supplementary Information for "A CUG-initiated CATSPERθ functions in the CatSper channel assembly and serves as a checkpoint for flagellar trafficking"

**This PDF file includes:**

**SI Materials and Methods**
**Fig. S1,** related to Fig. 1, *Tmem249* coevolved with other CatSper components and expressed specifically in testis.
**Fig. S2,** related to Fig. 2, Comparison of CATSPERθ homologs in amniotes.
**Fig. S3,** related to Fig. 2, Conservation of CTG initiated CATSPERθ in different species.
**Fig. S4,** related to Fig. 3, Generation of *Catsperq* knockout mice by CRISPR/Cas9.
**Fig. S5,** related to Fig. 3, Comparison of sperm motility and CatSper subunits expression in *wt* and *Catsperq*$^{-/-}$ sperm.
**Fig. S6,** related to Fig. 5, CATSPERθ in the connecting interface of CatSper dimer.
**Fig. S7,** related to Fig. 4 and Fig. 5, Processing of CATSPERθ during sperm capacitation and a working model of CATSPERθ in the assembly of the CatSper complex.

Movies S1-S6

### SI Materials and Methods

**Histological analysis.** Testes were collected from adult mice and fixed in Bouin's fluid (Polysciences) at 4°C overnight. Fixed samples were dehydrated by increasing ethanol concentrations and embedded with paraffin. Paraffin sections (5 μm) were rehydrated and treated with 1% periodic acid solution (Wako) for 15 min, followed by treatment with Schiff's reagent (Wako) for 20 min, and then Mayer's hematoxylin solution (Wako) for 5 min. After dehydration with ethanol, these slides were mounted with Permount (Fisher Scientific) and observed with a BX-53 microscope (Olympus).

**RNA extraction, cDNA synthesis, and RT-PCR.** Total RNA was extracted from adult *wt*, homozygous *Catsperq*$^{-/-}$, *Catsper1*$^{-/-}$ and *Catsperd*$^{-/-}$ male testes using RNeasy Mini kit (QIAGEN). 500 ng of the extracted RNA was used for cDNA synthesis using iScript cDNA Synthesis kit (Bio-Rad) according to manufacturer's instruction. cDNAs were subjected to quantitative PCR (qPCR) using iTaq Universal SYBR Green Supermix (Bio-Rad). Primers, forward (5'-GCCAATCAGCTCAAGAATAACAG-3'), reverse, (5'-AGGAAATGAAGACAAGGCAGA-3'), were used for qPCR. TBP was used as a reference gene to normalize transcript levels by ddCt method.

**Recombinant protein expression in mammalian cells and immunoprecipitation.** Different mouse *Catsperq* ORF and truncations were subcloned into phCMV3 backbone. CATSPER4 and truncations were subcloned into pcDNA3.1 backbone. HEK293T cells were transiently transfected with indicated plasmids with Lipofectamine 2000 (Invitrogen), following the manufacturer's instruction. Transfected cells were used for co-IP experiments. Briefly, after transfection for 18-20h, cells were lysed with 1% Triton X-100 in PBS containing EDTA-free protease inhibitor cocktail (Roche) by rocking at 4°C for 1h and centrifuged at 18,000 x g for 30 min at 4°C. Solubilized proteins in the supernatant were mixed with anti-HA magnetic beads (Pierce, 88836) at 4°C overnight and washed with 0.1% Triton PBS for three times. Co-IP products were eluted with 2x LDS sampling buffer supplemented with 50 mM dithiothreitol (DTT) and denatured at 75°C for 10 min. Primary antibodies used for the western blotting were rabbit monoclonal anti-HA (CST, #3724), and rabbit polyclonal anti-Flag (CST, #14793). For secondary antibodies, anti-rabbit IgG-HRP (Jackson ImmunoResearch) were used.

**Preparation of whole sperm lysate and solubilized protein extracts.** Mouse epididymal spermatozoa washed in PBS were directly lysed in 2x SDS sample buffer. Then the whole sperm lysates were centrifuged at 18,000 x g, 4°C for 10 min. After adjusting DTT to 50 mM, supernatant was denatured at 75°C for 10 min before loading to gel.

**Mouse sperm preparation and in vitro capacitation.** Epididymal spermatozoa from adult male mice were collected by swim-out from caudal epididymis in M2 medium (EMD Millipore, MR-015-D). Collected sperm were incubated in human tubular fluid HTF medium (EMD Millipore, MR-070-D) at $2 \times 10^6$ cells/ml concentration to induce capacitation at 37°C, 5% $CO_2$ for 90 min.

**Structured illumination microscopy.** Structured illumination microscopy (SIM) imaging was performed with Zeiss LSM710 Elyra P1 using alpha Plan-APO 100X/1.46 oil objective lens. Samples were prepared as described in Sperm Immunocytochemistry. z stack images were acquired with 200 nm intervals and each section was taken using 5 grid rotations with a 51 nm SIM grating period and a laser at 561 nm wavelength. Raw images were processed and rendered using Zen 2012 SP2 software (Carl Zeiss).

**Motility analysis.** Cauda epididymal spermatozoa were dispersed in a drop of TYH medium and incubated for 10 min and 2 h at 37°C under 5% $CO_2$. Spermatozoa were collected from the top of the drop and analyzed with CEROS II (software version 1.11.9; Hamilton Thorne Biosciences) sperm analysis system. Sperm motility (%) and progressive motility (%) were quantified, and motion parameters including straight line velocity (VSL), average path velocity (VAP) and curvilinear velocity (VCL) were measured. Spermatozoa were considered progressively motile when VSL/VAP ≥ 0.8 and VAP ≥ 50 μm/s.

**Antibodies and Reagents.** Rabbit polyclonal antibodies specific to mouse CATSPER1, 3, 4, β, δ, ζ and EFCAB9 were described previously (1-5). To produce antibody recognizing mouse CATSPERθ, peptide corresponding to mouse CATSPERθ (229-242, TQVYTKSSVNDLDV) was synthesized and conjugated to KLH carrier protein (GenScript). Antisera from the immunized rabbits were affinity-purified using the peptide immobilized SulfoLink Coupling Resin (Thermo, 20401). Anti-HA (CST, #3724), anti-Flag (CST, #14793), anti-acetylated Tubulin (Sigma, T7451).

**AlphaFold-Based CATSPERθ structure prediction, multiple alignment, and phylogenetic reconstruction.** The structure of CATSPERθ (242 amino acids) is generated with AlphaFold v2.09 on the Yale High Performance Cluster. Representative CATSPERθ sequences were aligned using Mafft at default settings. The MSA figure was created with ESPript 3.0 (https://espript.ibcp.fr/ESPript/cgi-bin/ESPript.cgi). Maximum likelihood tree was constructed with IQ-TREE using the automatically selected substitution model LG+I+G4. The tree figure was created with FigTree v1.4.3 (http://tree.bio.ed.ac.uk/software/figtree/).

**Fitting of CatSper cryo-EM and cryo-ET**. The single particle cryo-EM structure and corresponding atomic model of isolated CatSper complex (Lin et al) were fitted as a rigid body into the cryo-ET structure of the in situ CatSper complex (Zhao, et al) using the “fit-in-map” functionality in UCSF chimera. The protein density map of CATSPERθ was isolated by “volume zone” functionality in Chimera X-1.3.

**Motility correlative Imaging.** Live sperm motility was recorded as in **Flagellar waveform analysis**, but with photo-etched coverslips (Electron Microscopy Sciences #72265-12). Immediately after video recording, the coverslip was used for immunostaining CATSPERθ using the same method as in **Sperm immunocytochemistry**. Images photographed at the focus of the sperm and the focus of the engraved coordinates on the coverslip from the same field were merged. The cells on the merged images were backtracked to the video to identify the sperm motility.

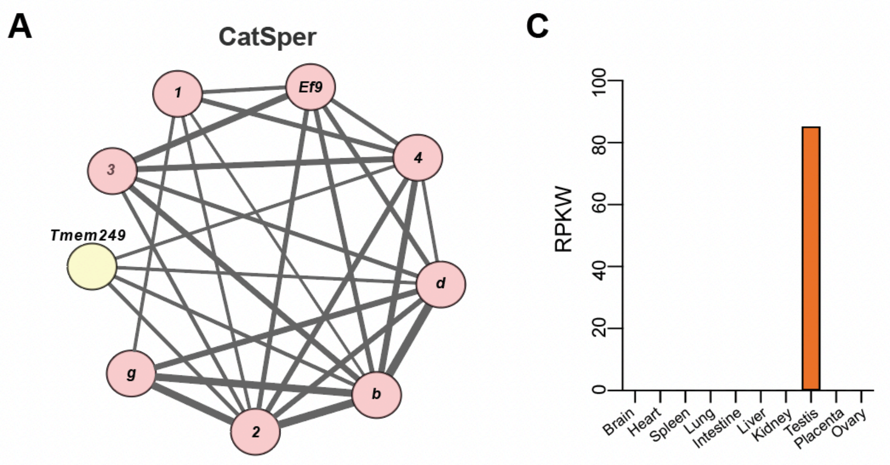


**B**

| OrthoDB ID1 (gene) | OrthoDB ID2 (gene) | T1 | T2 | C | D | Score | P_adj. |
|---|---|---|---|---|---|---|---|
| 1380438at2759 *(Tmem249)* | 846266at2759 ( ***Catssperb*** ) | 34 | 27 | 14 | 1 | 0.271 | 9.2E-06 |
| 1380438at2759 *(Tmem249)* | 580290at2759 ( ***Catsper2*** ) | 34 | 39 | 15 | 0 | 0.259 | 2.8E-06 |
| 1380438at2759 *(Tmem249)* | 213907at2759 ( ***Catsperd*** ) | 34 | 35 | 14 | 0 | 0.255 | 1.6E-05 |
| 1380438at2759 *(Tmem249)* | 1270192at2759 ( ***Catsper4*** ) | 34 | 36 | 14 | 0 | 0.250 | 2.6E-05 |
| 1380438at2759 *(Tmem249)* | 1039290at2759 (N.A.) | 34 | 24 | 11 | 0 | 0.234 | 3.5E-03 |
| 1380438at2759 *(Tmem249)* | 1269955at2759 ( ***Catsper3*** ) | 34 | 40 | 14 | 1 | 0.213 | 4.5E-03 |
| 1380438at2759 *(Tmem249)* | 110808at2759 ( ***Catsperg*** ) | 34 | 35 | 13 | 1 | 0.211 | 1.9E-02 |
| 1380438at2759 *(Tmem249)* | 313028at2759 *(Wdr97)* | 34 | 43 | 13 | 0 | 0.203 | 1.3E-02 |
| 1380438at2759 *(Tmem249)* | 679293at2759 *(Tmem232)* | 34 | 17 | 9 | 0 | 0.214 | 9.6E-02 |
| 1380438at2759 *(Tmem249)* | 1306120at2759 ( ***Efcab9*** ) | 34 | 38 | 12 | 0 | 0.200 | 5.7E-02 |

**Fig. S1.** *Tmem249* has coevolved with other CatSper components and is expressed specifically in testis. (*A*) Coevolutionary analysis of CatSper network obtained by automatic clustering (Markov cluster, MCL) of significant pairwise associations (p_adj <0.001; Fisher exact test with Bonferroni correction) among 60675 OrthoDB gene groups in 1256 eukaryotic organisms. Known CatSper components are colored pink; *Tmem249* is colored yellow. Edge thickness is proportional to the p-value significance. (*B*) The ten most significant coevolutionary associations of *Tmem249* in the complete dataset (known CatSper components highlighted in pink) showing the total number of transitions (absence/presence) of gene1 (T1) and gene2 (T2), the number of concordant (C) and discordant (D) transitions, the score [k/(T1+T2-k), where k=C-D], and the adjusted p-value. (*C*) Expression profile of *Tmem249* mRNA among various tissues. Data sets are generated according to the Mouse ENCODE project. Data source: https://www.ncbi.nlm.nih.gov/gene/666504/?report=expression.

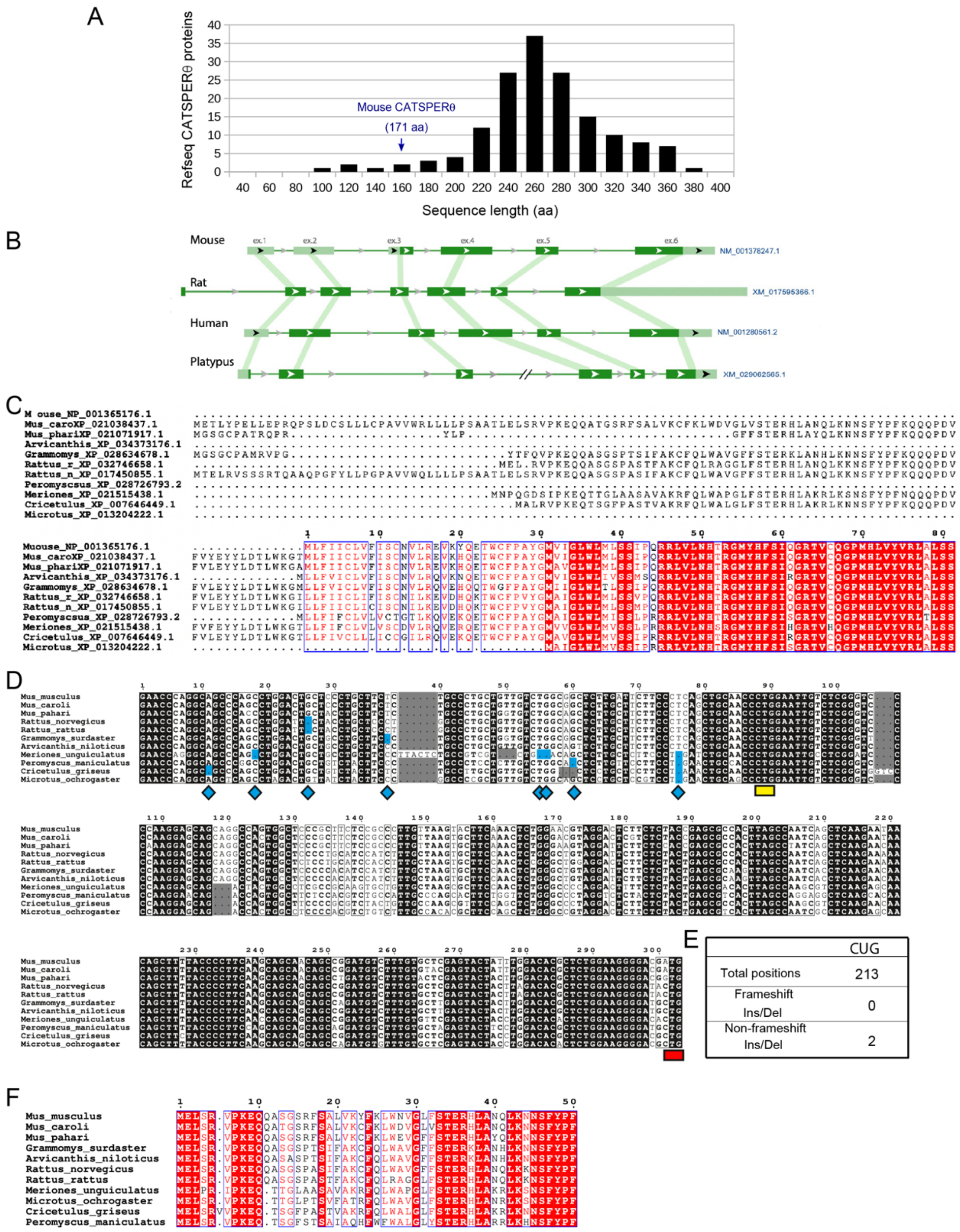


**Fig. S2**. Comparison of CATSPERθ homologs in amniotes. (*A*) Protein length distribution of CATSPERθ (available ref-seq proteins) from 162 different amniotes. (*B*) Comparison of the *Catsperq* genomic structure in different mammals. The exon/intron structure is according to the NCBI gene record; the corresponding mRNA reference sequences are indicated by database accession numbers. Exons are represented by horizontal segments with the coding sequence in dark green. Homologous exons in different species are connected by vertical segments. (*C*) Multiple CATSPERθ protein sequences from the original NCBI database are aligned with their N-terminal region. The number above the alignment indicate the amino acid number of protein CATSPERθ from *Mus musculus* initiated by ATG. (*D*) Multiple sequence alignment of rodent *Catsperq* 5'-UTRs; frameshift mutations are highlighted in blue, non-frameshift mutations are highlighted in gray, the candidate CTG start codon is indicated by yellow bar. The red bar indicates the position of the original NCBI assigned ATG start codon in the mouse sequence. (*E*) Occurrence of frameshift and non-frameshift mutations in the alignment of rodent *Catsperq* 5'-UTRs. (*F*) Multiple protein sequence alignment of the rodent CATSPERθ N-terminal region translated from CTG. In (*C* and *F*) columns with identical residues are highlighted in red and conserved residues are boxed in blue.

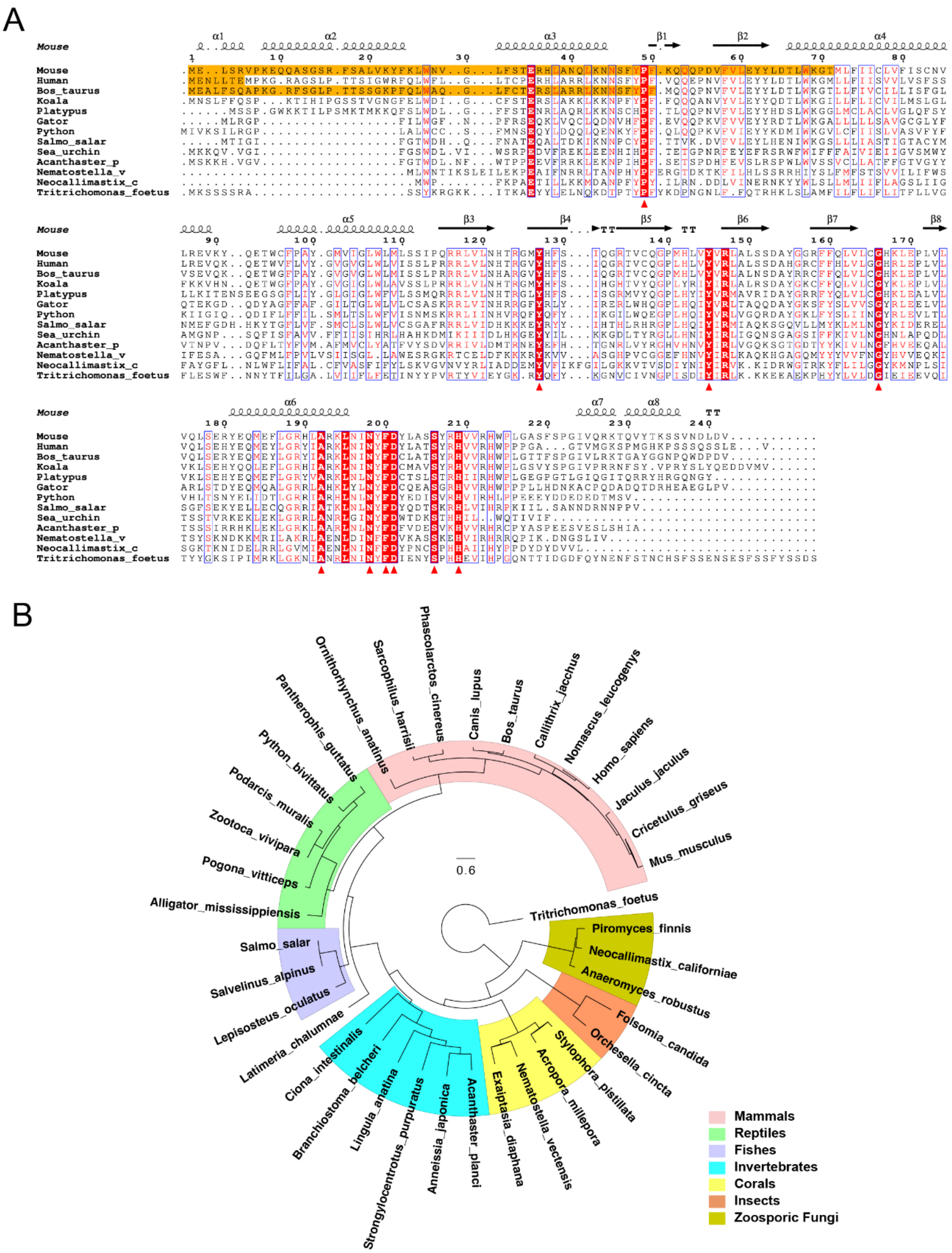
A
B
Phascolarctos_cinereus
Sarcophilus_harrisii
Ornithorhynchus_anatinus
Pantherophis_guttatus
Python_bivittatus
Podarcis_muralis
Zootoca_vivipara
Pogona_vitticeps
Alligator_mississippiensis
Salmo_salar
Salvelinus_alpinus
Lepisosteus_oculatus
Latimeria_chalumnae
Ciona_intestinalis
Branchiostoma_belcheri
Lingula_anatina
Strongylocentrotus_purpuratus
Anneissia_japonica
Acanthaster_planci
Exaiptasia_diaphana
Nematostella_vectensis
Acropora_millepora
Stylophora_pistillata
Orchesella_cincta
Folsomia_candida
Anaeromyces_robustus
Neocallimastix_californiae
Piromyces_finnis
Tritrichomonas_foetus
Mus_musculus
Cricetulus_griseus
Jaculus_jaculus
Homo_sapiens
Nomascus_leucogenys
Callithrix_jacchus
Bos_taurus
Canis_lupus
0.6
Mammals
Reptiles
Fishes
Invertebrates
Corals
Insects
Zoosporic Fungi

C

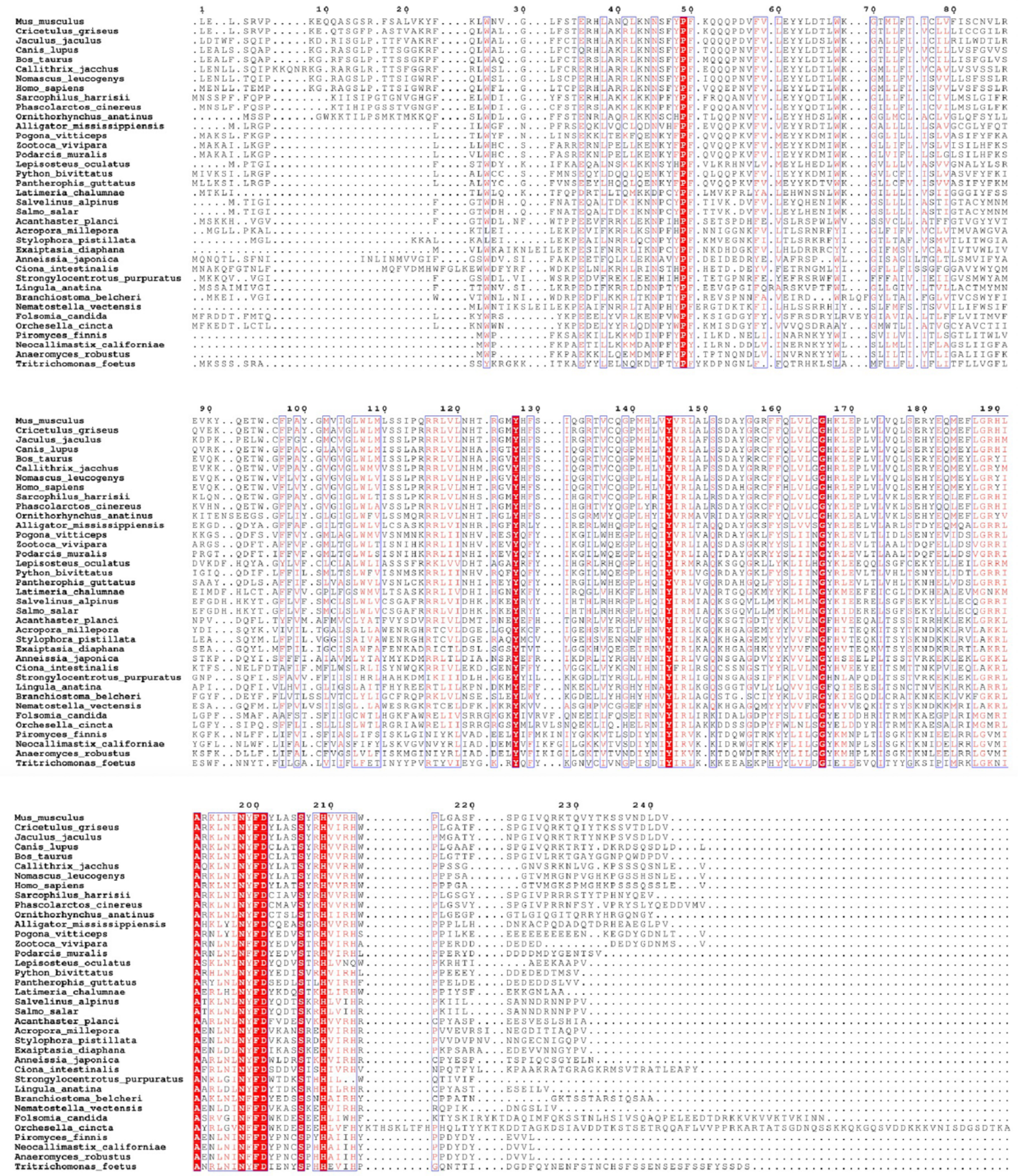

**Fig. S3.** Conservation of CTG initiated CATSPERθ in different species. (*A* and *C*) Multiple alignment of representative CATSPERθ protein sequences. Columns with identical residues are highlighted in red and conserved residues are boxed in blue; residues that are conserved in a full alignment of 37 sequences (*C*) are indicated by arrowheads. Mammalian coding regions differing from database sequences due to translation from the putative CTG starting codon are shaded in orange. (*B*) Maximum likelihood tree of CATSPERθ sequences showing phylogenetic relations in agreement with organism phylogeny. Different taxonomic groups are boxed in different colors. The polarized circular tree was rooted using *Tritrichomonas foetus* as an outgroup.

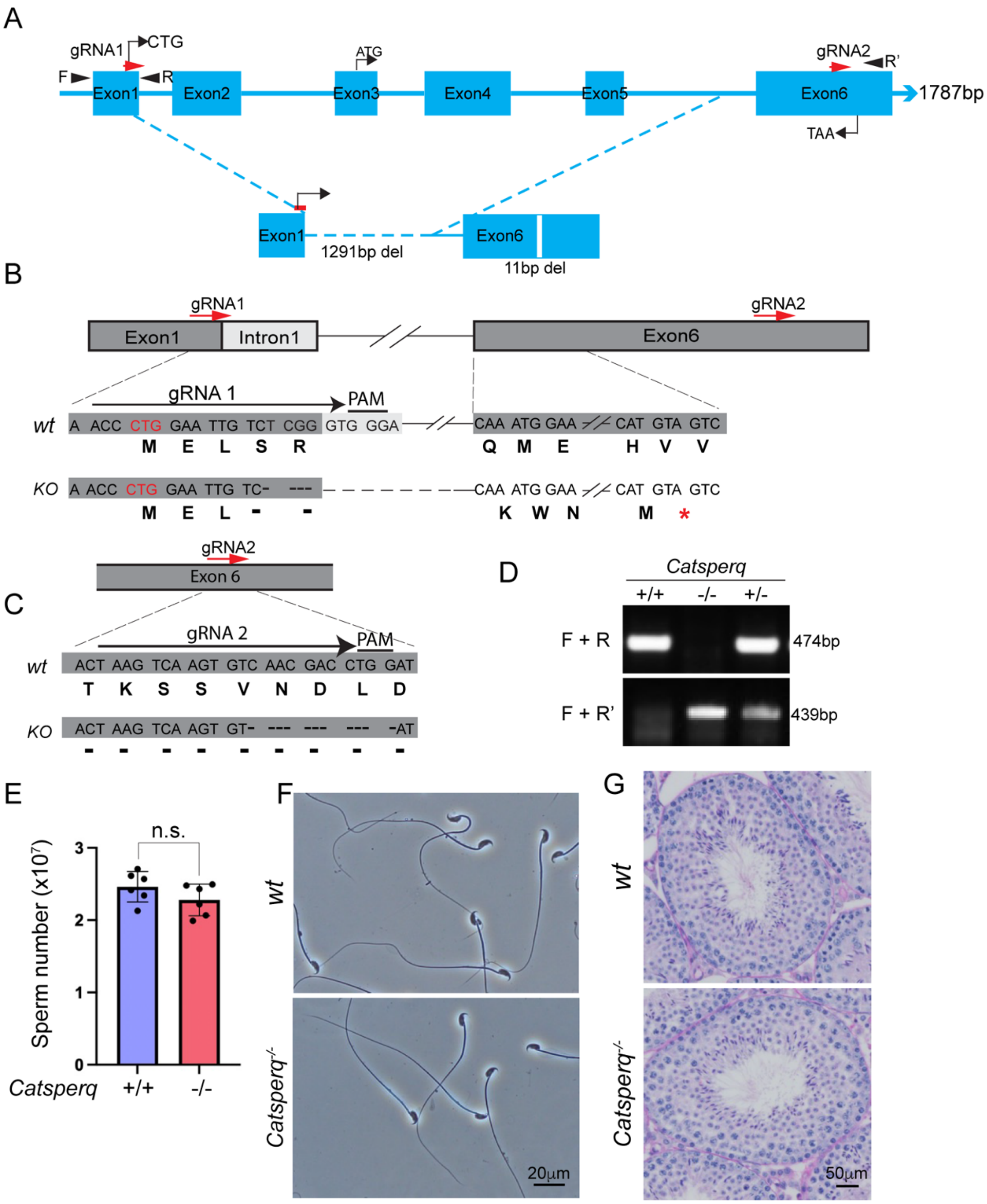


**Fig. S4.** Generation of *Catsperq* knockout mice by CRISPR/Cas9. (*A*) Schematic presentation of mouse *Catsperq* genomic structure showing *wt* and CRISPR/Cas9-induced knockout. Red arrows indicate the target location of two gRNAs used for CRISPR/Cas9. The locations of the genotyping primers are indicated as F, R and R'. (*B-C*) Knockout allele with 1291 bp and 11bp deletion at genomic region encoding CATSPERθ. The deletions are expected to cause out-of-frame mutations, resulting in early termination of protein translation from the CTG start codon. (*D*) Genomic DNA PCR for genotyping of *Catsperq* knockout mice with the primer pairs marked in (*A*). (*E*) Epididymal sperm count from littermates at ages 3-6 months. *wt* (blue) versus *Catsperq*$^{-/-}$ (red) sperm. Data is represented as mean SEM. (*F*) Sperm morphology. (*G*) Histological analysis of testes from *wt* and *Catsperq*$^{-/-}$ mice.

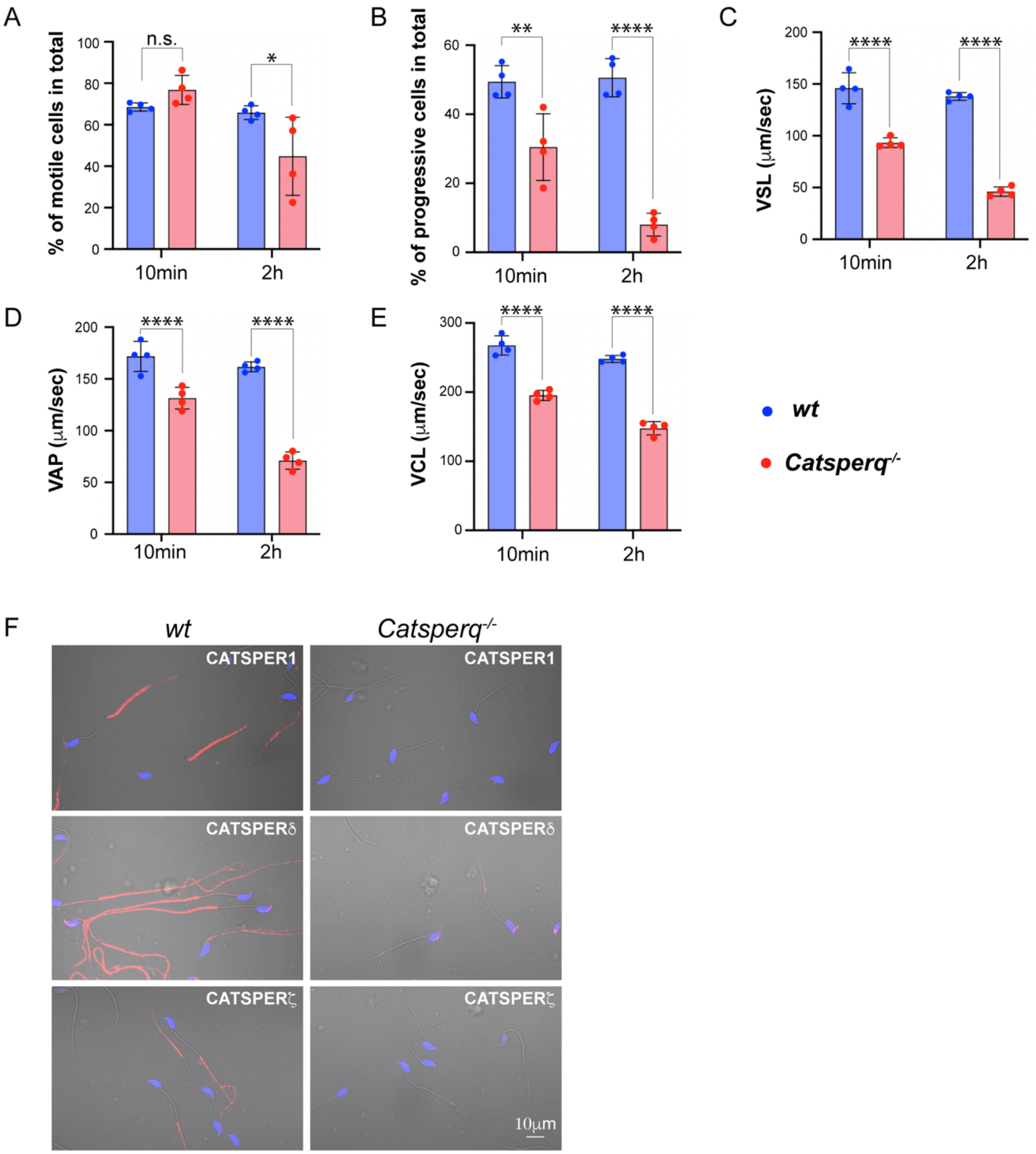


**Fig. S5.** Comparison of sperm motility and CatSper subunits expression in *wt* and *Catsperq*$^{-/-}$ sperm. The motility parameters of *wt* (blue bars) and *Catsperq*$^{-/-}$ (pink bars) sperm were measured from sperm after 10 min and 2h incubation with TYH capacitation media by computer-assisted sperm analysis (CASA). (*A*) Total motility. (*B*) Progressive motility. (*C*) Straight linear velocity (VSL). (*D*) Average path velocity (VAP). (*E*) Curve linear velocity (VCL). *Catsperq*$^{-/-}$ sperm shows comprised motility endurance over time together with defective hyperactivation. *P<0.5, **P<0.01, ****P<0.0001. Data are presented as mean $\pm$ SEM, n=4. (*F*) Absent or barely detectable protein levels of other CatSper subunits in *Catsperq*$^{-/-}$ sperm by immunocytochemistry.

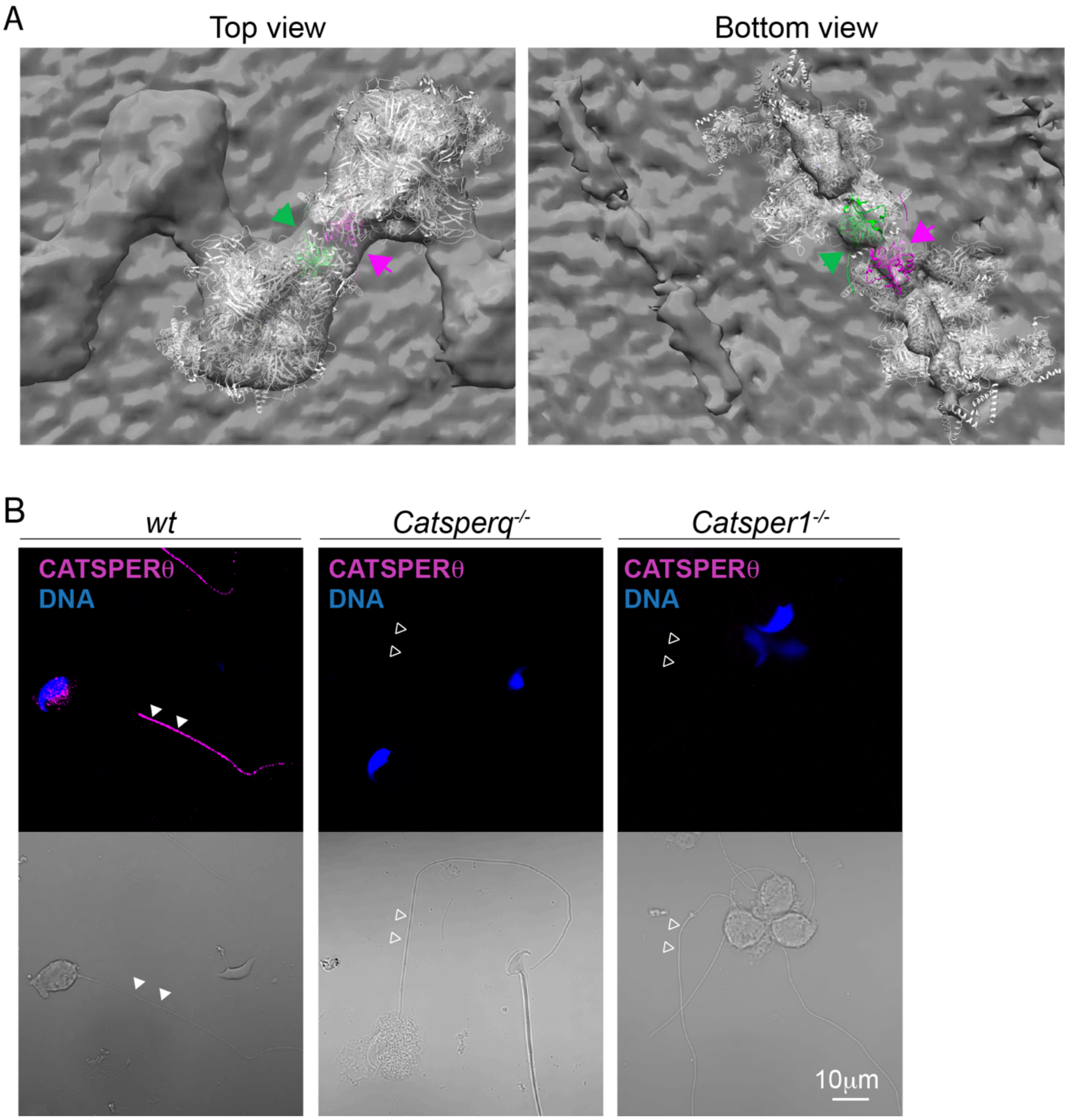


**Fig. S6.** CATSPERθ in the connecting interface of CatSper dimer. (*A*-*B*) Top view (*left*) and bottom view (*right*) of two CatSper complex modified from (6) in ribbon presentation docked into the averaged CatSper cryo-ET map (7). Two CATSPERθ from each monomeric CatSper complex were highlighted with green and magenta, as the green and magenta arrows indicate. (*B*) Immunostaining of CATSPERθ in *wt* (*left*), *Catsperq*$^{-/-}$ (*middle*) and *Catsper1*$^{-/-}$ (*right*) developing spermatids (step8-10). The corresponding DIC images were shown (*lower*). Filled arrowheads indicate the presence of signal in the tail; empty arrowheads indicate the absence of the signal.

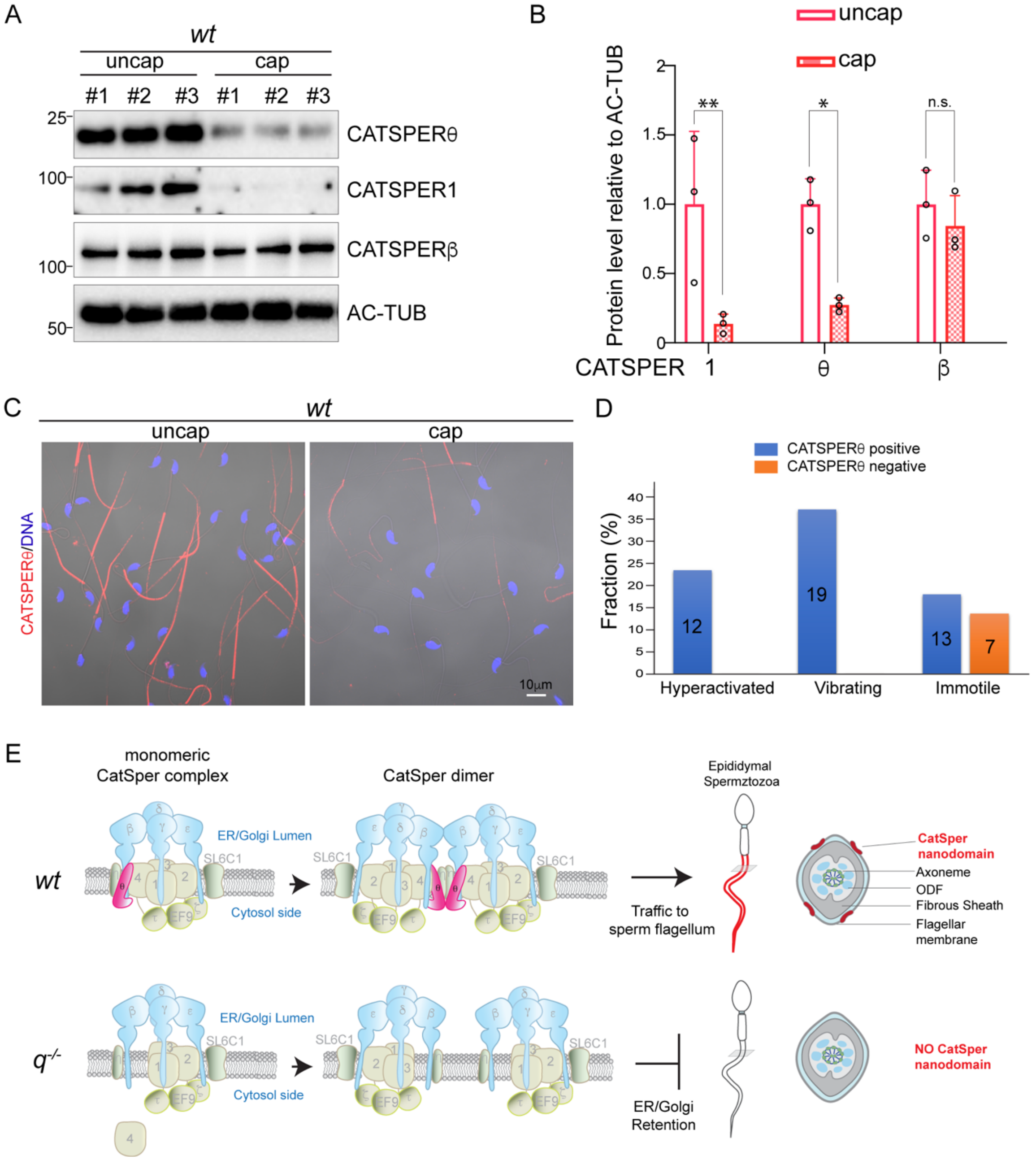


**Fig. S7**. Processing of CATSPERθ during sperm capacitation and a working model of CATSPERθ in the assembly of CatSper complex. (*A*) CATSPERθ, CATSPER1 and CATSPERβ protein levels are evaluated in *wt* sperm before and after 90 min capacitation by western blot. #1, #2 and #3 indicates sperm samples collected from three mice, respectively. (*B*) Quantitative protein level relative to AC-TUB in (*A*). *P<0.5, **P<0.01, Data are presented as mean ± SEM, n=3. (*C*) Representative immunofluorescence images of CATSPERθ (red) before and after capacitation. (*D*) CATSPERθ positive or negative fraction within sperm with different motility behavior. The numbers represent the number of sperm counted for each group. (*E*) Working model of CATSPERθ in CatSper complex assembly. In *wt* spermatid, a single CatSper complex unit can assemble normally and dimerize with another properly assembled CatSper complex. CatSper complex can traffic to sperm flagellum and form the 4 linear nanodomains. While without CATSPERθ ($q^{-/-}$), CATSPER4 is not stable in the CatSper complex unit. And the absence of CATSPERθ in CatSper complex abolish/destabilize the CatSper dimer formation, thus preventing them from trafficking to sperm flagellum. SL6C1, SLCO6C1; EF9, EFCAB9.

**SI References**


1. D. Ren *et al.*, A sperm ion channel required for sperm motility and male fertility. *Nature* **413**, 603-609 (2001).
2. J. J. Chung, B. Navarro, G. Krapivinsky, L. Krapivinsky, D. E. Clapham, A novel gene required for male fertility and functional CATSPER channel formation in spermatozoa. *Nat Commun* **2**, 153 (2011).
3. H. Qi *et al.*, All four CatSper ion channel proteins are required for male fertility and sperm cell hyperactivated motility. *Proc Natl Acad Sci U S A* **104**, 1219-1223 (2007).
4. J. J. Chung *et al.*, CatSperzeta regulates the structural continuity of sperm Ca2+ signaling domains and is required for normal fertility. *Elife* **6** (2017).
5. J. Y. Hwang *et al.*, Dual Sensing of Physiologic pH and Calcium by EFCAB9 Regulates Sperm Motility. *Cell* **177**, 1480-1494 e1419 (2019).
6. S. Lin, M. Ke, Y. Zhang, Z. Yan, J. Wu, Structure of a mammalian sperm cation channel complex. *Nature* **595**, 746-750 (2021).
7. Y. Zhao *et al.*, 3D structure and in situ arrangements of CatSper channel in the sperm flagellum. *Nat Commun* **13**, 3439 (2022).
